# Targeting the NuRD Component, CHD4, Impairs Foxp3+ Treg Cell Production and Function and Promotes Anti-Tumor Immunity

**DOI:** 10.64898/2026.07.30.741771

**Authors:** Yan Xiong, Liqing Wang, Martina Minisini, Fanhua Kong, Eros di Giorgio, Tatiana Akimova, Shane R. Horman, Ivan Babic, Elmar Nurmemmedov, Wayne W. Hancock

**Author notes:** These authors contributed equally: Elmar Nurmemmedov and Wayne W. Hancock.

## Abstract

Little is known about why Foxp3⁺ regulatory T (Treg) cells require at least three HDAC1/HDAC2-containing chromatin-remodeling complexes (NuRD, Sin3 and CoREST), or whether selective disruption of these complexes can be exploited to enhance antitumor immunity. Here, we investigated the role of chromodomain helicase DNA-binding protein 4 (CHD4), the ATP-dependent remodeling subunit of the NuRD complex, in Treg biology. Conditional deletion of Chd4 in Foxp3⁺ Tregs resulted in severe systemic autoimmunity and early lethality, accompanied by reduced Foxp3 expression, impaired Treg suppressive function, and loss of Treg lineage stability. Transcriptomic analyses demonstrated that CHD4 deficiency closely phenocopied Hdac2 deletion, whereas quantitative proteomic analyses revealed that CHD4 assembles into highly conserved NuRD complexes in both Treg and conventional CD4⁺ T cells. These findings indicate that the selective dependence of Tregs on CHD4 does not arise from the formation of lineage-specific protein complexes but rather from the unique epigenetic program maintained by CHD4-containing chromatin-remodeling complexes that is required for Treg differentiation and stability. Using a novel cellular target-engagement platform, we identified CH41, a potent small-molecule inhibitor of CHD4 that recapitulated the effects of genetic CHD4 ablation on Treg function. Pharmacological inhibition of CHD4 impaired intratumoral Treg accumulation and function and significantly inhibited the growth of lung and hepatocellular carcinomas in immunocompetent, but not immunodeficient, mice, without inducing systemic autoimmunity. Collectively, our findings identify CHD4 as a critical epigenetic regulator of Treg lineage stability and establish pharmacological targeting of the CHD4/NuRD axis as a promising strategy to selectively disrupt tumor-associated Tregs and enhance antitumor immunity.

## INTRODUCTION

Foxp3^+^ CD4^+^ Treg cells are essential for maintaining immune tolerance and preventing autoimmune diseases (1, 2), though these cells also dampen host antitumor immunity, decreasing the efficacy of tumor immune surveillance (3). The transcription factor Foxp3 has a critical role in the differentiation and function of Tregs (4, 5), and knockdown or mutations of Foxp3 attenuate the immunosuppressive capacity of Treg cells (6, 7). Thus, depletion of Foxp3+ CD4+ Tregs results in severe autoimmunity in otherwise normal animals but can be reversed by Treg reconstitution (8, 9). Recent successes with checkpoint inhibitor therapies in the treatment of various cancers have rekindled interest in immunotherapy, but despite a major contribution of Foxp3+ Tregs to establishing and maintaining an immunosuppressive tumor microenvironment, there are few options to selectively target Foxp3+ Tregs and promote antitumor immunity (10, 11).

Chromodomain helicase DNA-binding 4 (CHD4), an ATP-dependent chromatin remodeling helicase also known as Mi-2β, is a protein within the NuRD complex that also includes histone deacetylases HDAC1/HDAC2, RBBP4/RBBP7 (also known as RbAp48/46), GATAD2A/GATAD2B, MTA1/MTA2/MTA3, the mCpG-binding domain proteins MBD2/MBD3, and the histone demethylase KDM1A (12). These components perform functions such as histone deacetylation, ATP-dependent chromatin remodeling, histone chaperoning, CpG binding, DNA-binding, and transcriptional regulation (13). CHD4 is a pleiotropic factor promoting DNA damage repair, cell cycle transition, and the development of immune and neuronal cells (14, 15). While CHD4 helps regulate T and B lymphocyte development and differentiation (14, 16, 17), its role in cancer is less well understood, though mutations in CHD4 can promote tumorigenesis, particularly in endometrial cancer, by enhancing cancer stem cell (CSC) characteristics and activating oncogenic signaling pathways (18). Genetic alterations in CHD4 reduce protein stability and function, effectively mimicking CHD4 loss-of-function, and commonly occur in hepatocellular, breast, ovarian, colorectal and non-small cell lung cancers (19).

A prior large-scale mass-spectrometric survey of proteins complexed with Foxp3 in transfected cells demonstrated Foxp3 forms multiprotein complexes of 400–800 kDa or larger and identified 361 associated proteins, including components of the NuRD, SWI/SNF, NCoR and other complexes (20). However, the role of CHD4 in the regulation of Foxp3+ Tregs has not been previously studied. Accordingly, we conditionally deleted CHD4 in Treg cells and tested the effects of recently developed CHD4 inhibitors in murine models. We found that gene deletion or pharmacologic inhibition disrupted Foxp3-dependent recruitment of the CHD4 complex to the promoter of IL2, impaired Treg function, and enhanced antitumor immunity, adding new insights into the functions of CHD4 and its potential therapeutic targeting in cancer.

## RESULTS

### CHD4 deletion led to decreased Foxp3 expression and lethal autoimmunity

Foxp3 has a central role in maintaining Treg stability and function and forms multiprotein nuclear complexes (≥400–800 kDa) that contain various transcription factors and corepressor proteins (20). However, the functions of these evolutionarily highly conserved nuclear complexes in Treg cells are largely unexplored. We began by assessing whether CHD4 could bind to Foxp3. In 293T cells transfected with tagged constructs, immunoprecipitation of CHD4 resulted in co-precipitation of Foxp3 (55 kDa) and Tip60 (60 kDa) (Fig. 1a). We have previously shown that immunoprecipitation of Foxp3 leads to co-precipitation of Tip60, which is important for Foxp3 acetylation, dimerization, and Treg function (21). Hence, the association of CHD4 with Foxp3 and Tip60 may contribute to control of Treg functions and immune homeostasis. To further explore this point, we conditionally deleted CHD4 in Foxp3+ Treg cells by crossing CHD4^fl/fl^ and Foxp3^YFP/Cre^ mice (Fig. 1b). CHD4^fl/fl^Foxp3^YFP/Cre^ (hereafter CHD4^-/-^) mice were born at expected Mendelian ratios but died by 3-4 weeks of life in conjunction with severe weight loss, dermatitis, lymphadenopathy and splenomegaly (Fig. S1a), mononuclear infiltrates in lung and liver (Fig. S1b), and disrupted hematopoiesis with markedly decreased RBC, hemoglobin, hematocrit and platelet levels (Fig. S1c). Histologic findings are summarized in Suppl. Table 1, and sera of CHD4^-/-^ mice contained multiple autoantibodies (Fig. S2, Suppl Table 2). CHD4^-/-^ Tregs had decreased expression of Foxp3 protein (Westerns & flow cytometry) (Fig. 1b-d), and the mean fluorescence intensity (MFI) of Foxp3 was also decreased in lymph nodes, spleen and thymus (Fig. 1d). Compared to littermates, CHD4^-/-^ mice showed increased T cell activation and production of proinflammatory cytokines (Fig. 1e, f), and the *in vitro* suppressive function of their Tregs was modestly impaired (P<0.05) compared with that of WT Tregs (Fig. 1g). Hence, CHD4 can associate with Foxp3, and under basal conditions, CHD4 deletion leads to decreased Foxp3 expression, decreased peripheral Treg numbers and Treg suppressive function, increased humoral response and the rapid development of lethal autoimmunity.

**Fig. 1.**
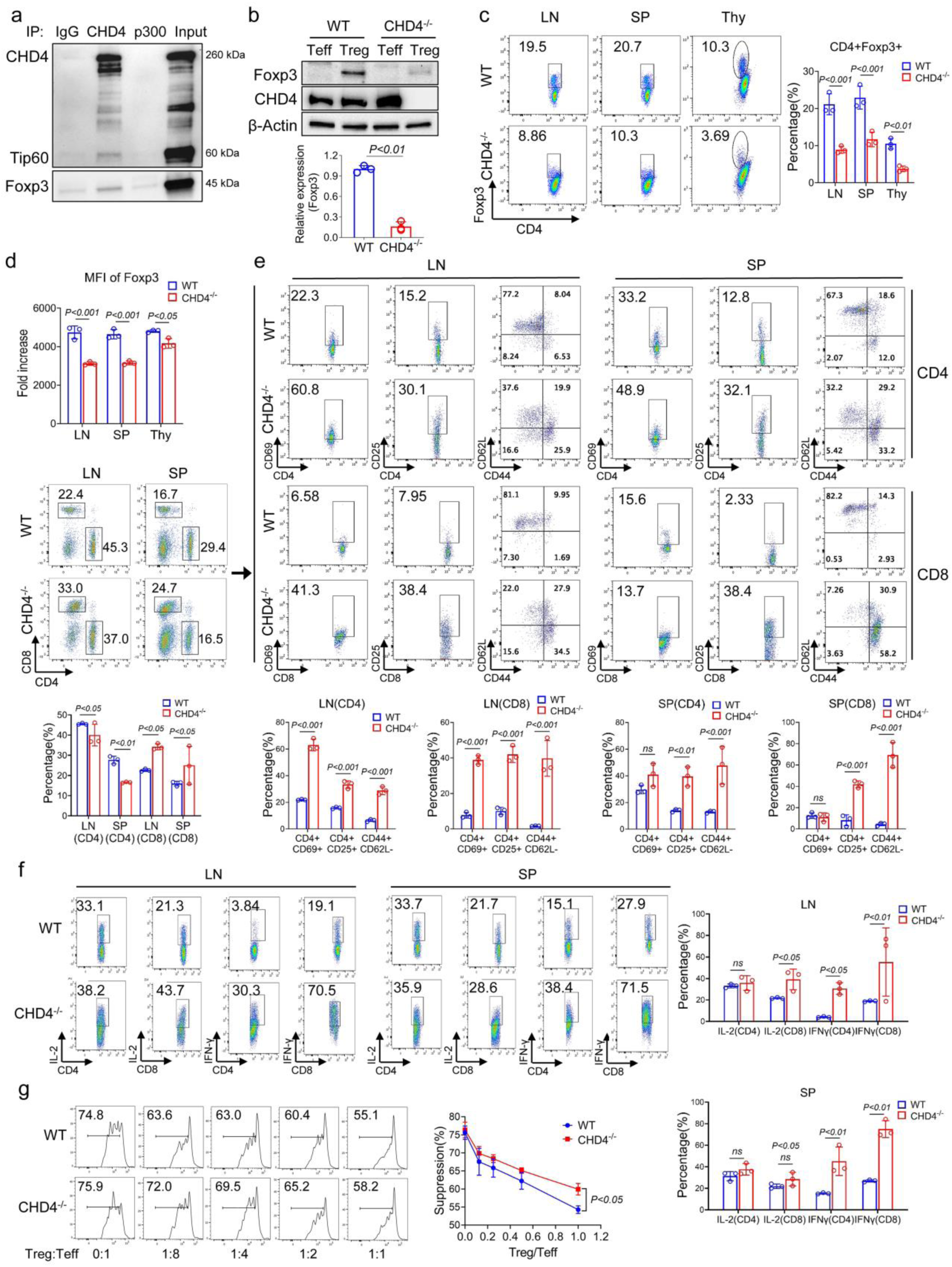
Effects of conditional CHD4 deletion in Foxp3^+^ Tregs. (**a**) HEK-293T cells were transfected with tagged constructs encoding Foxp3 (47 kDa), CHD4-Flag (250 kDa), Tip60-Flag (47 kDa) and p300-HA (300 kDa). Immunoprecipitation of CHD4 led to co-precipitation of Foxp3 and Tip60 but not p300 protein. (**b**) Western blots of CHD4 and Foxp3 expression in Treg and Teff cells from WT mice or mice with conditional deletion of CHD4 in their Tregs; β- actin was used as loading control. Representative of 2 independent experiments involving 3 mice/group at 20 d of life. P < 0.01 vs. WT control. (**c**) Percentages of CD4+Foxp3+ Tregs in lymph nodes, spleens and thymii of WT and CHD4^-/-^ mice, shown as representative plots (left) and with statistical analyses (right). (d) MFI value of Foxp3 analysis. (e) T cell activation markers and (f) T-cell production of IL-2 and IFN-γ when stimulated with PMA and ionomycin in the presence of GolgiStop for 5 h. Representative flow plots and overall data from 4-6 experiments. (**g**) Treg suppression assay using pooled Tregs from lymph nodes and spleens of WT and CHD4^-/-^ mice, with representative data shown, along with the percentage of proliferating cells in each panel. In (c - f) data are shown as mean ± SD, 6-8 mice/group at 20 d of life, ANOVA: *P<0.05; **P<0.01; ***P<0.001 vs. WT control.

### CHD4 deletion affected Treg gene expression

We undertook RNA-seq analyses of CHD4^−/−^and WT Tregs to assess CHD4-dependent global changes in gene expression. Compared with WT Tregs, 718 genes were upregulated, and 1399 genes were downregulated in CHD4 KO Tregs (>2-fold, P<0.05) (Fig. 2a). Gene set enrichment analysis revealed that genes downregulated following CHD4 deletion showed significant overlap with gene signatures downregulated in Foxp3-deficient Tregs and in Foxo1-deficient cells, the latter being a key transcription factor that primes Foxp3 expression. Conversely, CHD4-deficient Tregs displayed an inverse correlation with RORC-dependent gene signatures. These findings support a role for CHD4 in sustaining the FOXP3-dependent transcriptional program while antagonizing the RORγt-driven inflammatory program (Fig. 2b). Consistent with these findings, CHD4 deficiency resulted in the de-repression of cell cycle–associated genes, indicative of the loss of the differentiated Treg transcriptional program (Fig. 2c). Upregulation of various genes of interest, including IL-2 and apoptosis-linked genes such as A1a, Bag1, Bcl2, FasL and Mad2, was confirmed by quantitative PCR (qPCR) (Fig. 2d, e). Comparison of the transcriptomes of CHD4-deficient Tregs and B cells revealed that, while CHD4 regulates conserved pathways involved in fundamental cellular functions in both lineages, CHD4 loss in Tregs preferentially affected gene sets associated with Treg identity and suppressive function, highlighting a lineage-specific role for CHD4 in maintaining the Treg transcriptional program (Fig. 2f). Since CHD4 is part of the NuRD complex that includes HDAC1 and HDAC2, CHD4^-/-^ Tregs were subjected to GSEA comparison with Tregs lacking HDAC1 or HDAC2 (Fig. 2g). GSEA comparison revealed a substantial overlap between the transcriptional changes induced by CHD4 and HDAC2 deficiency, supporting the epigenetic role of the NuRD complex in maintaining Treg identity (Fig. 2h). Hence, impairment of Foxp3 functions in CHD4^-/-^ Tregs appear to stem from a deep epigenetic resetting controlled by the NuRD complex.

**Figure 2.**
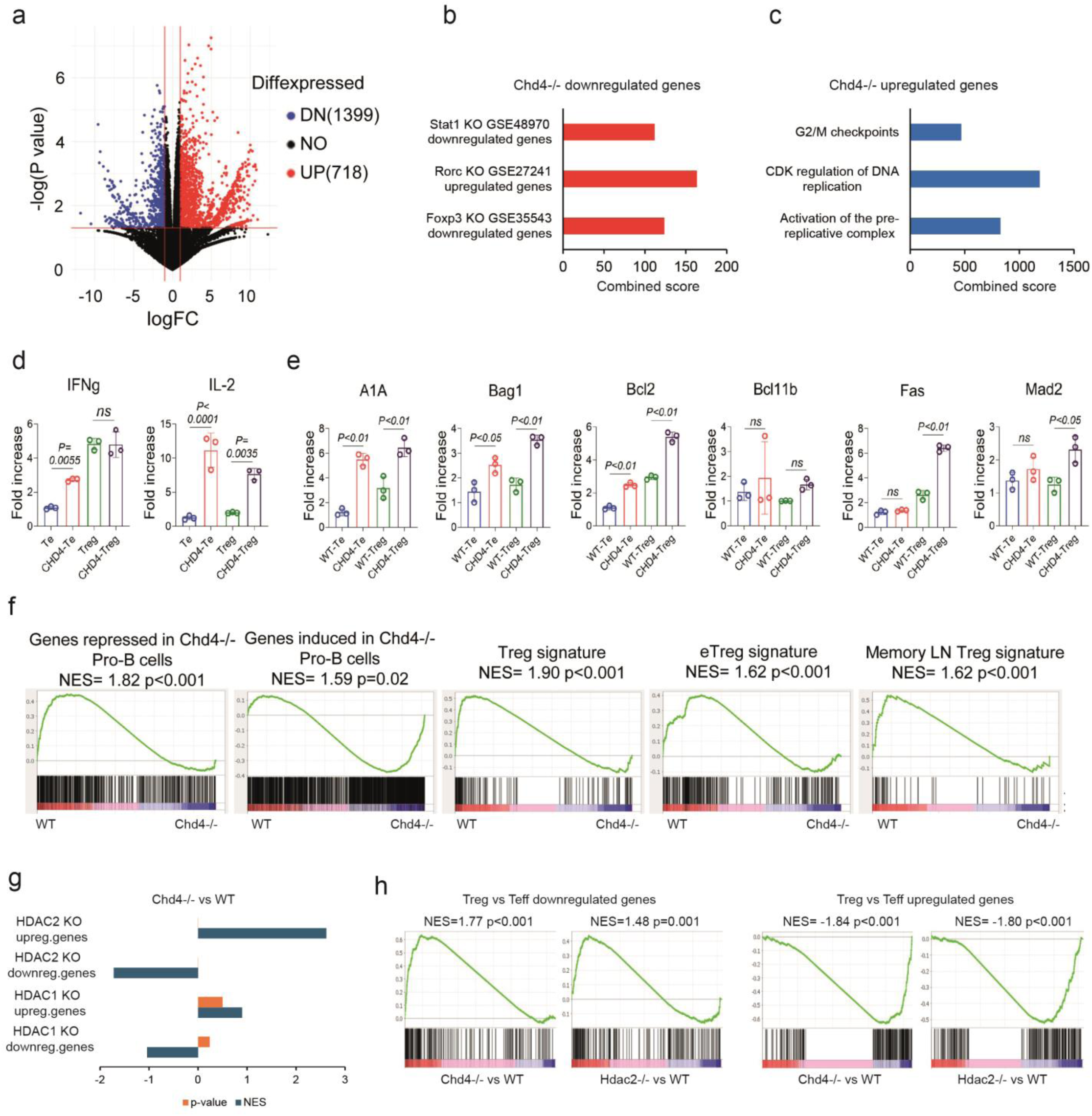
CHD4 deficiency disrupts the Treg transcriptional program while preserving the core epigenetic dependency on the NuRD complex. **(a)** Volcano plot showing differentially expressed genes in CHD4-deficient (Chd^-/-^) versus WT Treg cells. Genes significantly upregulated (red) and downregulated (blue) are indicated. **(b)** Enriched gene set enrichment analysis of genes downregulated in Chd4^−/−^ Tregs. Downregulated genes significantly overlapped with transcriptional signatures associated with Foxp3 deficiency, Stat1 deficiency, and genes induced by Rorc, supporting the loss of the canonical Treg transcriptional program. **(c)** Functional enrichment analysis of genes upregulated following CHD4 deletion. The most significantly enriched pathways were associated with cell-cycle progression, DNA replication, and activation of the pre-replicative complex, indicating loss of the differentiated Treg state. **(d, e)** Quantitative RT-PCR validation of the relative expression levels of selected cytokine genes (d) and genes involved in apoptosis and cell survival (e). **(f)** GSEA demonstrating enrichment of published transcriptional signatures in Chd4−/− Tregs in respect to wt Tregs. **(g)** Comparison of transcriptional changes induced by deletion of Chd4, Hdac1, or Hdac2. The transcriptional profile of Chd4-deficient Tregs showed a substantially greater similarity to that of Hdac2-deficient than Hdac1-deficient Tregs, supporting a predominant functional interaction between CHD4 and HDAC2 within the NuRD complex. **(h)** GSEA comparing transcriptional changes induced by Chd4 or Hdac2 deletion using published Treg-versus-Teff gene signatures. Data are presented as mean ± SD. Statistical significance was determined using two-tailed Student’s t-test or GSEA as indicated. NES, normalized enrichment score.

### CHD4 depletion affected Foxp3 expression

To determine whether CHD4 directly regulates Foxp3 expression through epigenetic mechanisms, we performed chromatin immunoprecipitation (ChIP) analysis to examine CHD4 occupancy at the Foxp3 locus. Chromatin was immunoprecipitated using an anti-CHD4 antibody, followed by qPCR analysis of the Foxp3 promoter and its conserved non-coding sequences (CNS0-CNS2) (Fig. 3a). ChIP-qPCR revealed specific CHD4 binding to the CNS1 intronic enhancer located downstream of the Foxp3 promoter in Treg cells (Fig. 3b). Consistent with a role for CHD4 in maintaining an active chromatin state, ChIP analysis of acetylated H3K27 demonstrated reduced enrichment at the Foxp3 promoter in CHD4-deficient Tregs compared with WT controls (Fig. 3c). Furthermore, in 293T cells, CHD4 significantly enhanced p65-mediated transcriptional activation of luciferase reporters driven by the Foxp3 promoter and the CNS1/CNS2 regulatory elements (Fig. 3d).

**Figure 3.**
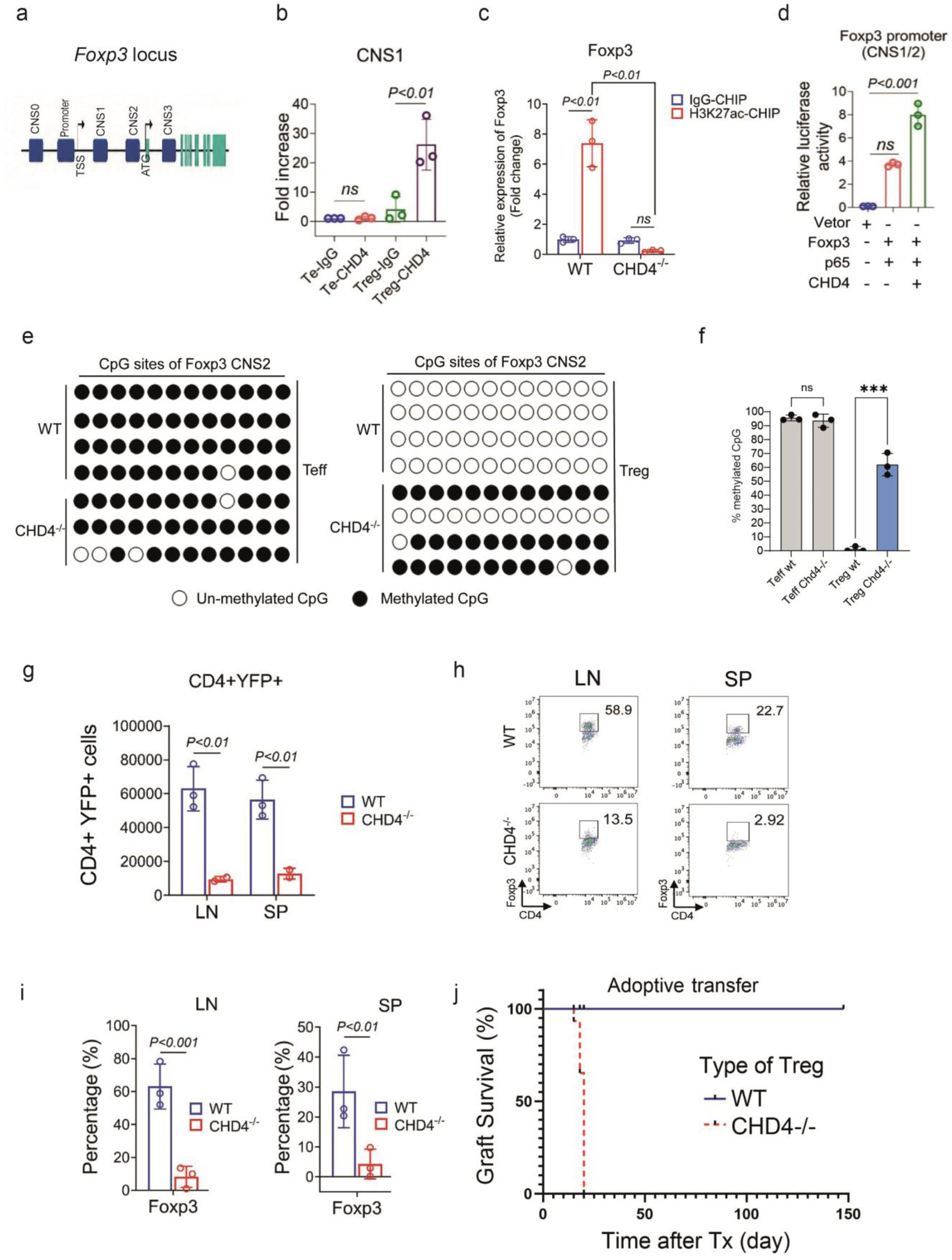
CHD4 maintains Foxp3 expression and Treg lineage stability through epigenetic regulation of the Foxp3 locus. (a) Schematic representation of the Foxp3 locus showing the promoter and the conserved non-coding sequences (CNS0–CNS3) analyzed in this study. (b) Chromatin immunoprecipitation (ChIP)-qPCR analysis of CHD4 binding to the Foxp3 locus. CHD4 was specifically enriched at the CNS1 enhancer in Treg cells but not in conventional CD4⁺ T cells (Teff). (c) ChIP analysis of H3K27 acetylation at the Foxp3 promoter. CHD4-deficient Tregs exhibited markedly reduced H3K27 acetylation compared with WT Tregs, consistent with decreased chromatin accessibility and reduced Foxp3 transcription. (d) Luciferase reporter assay performed in 293T cells demonstrating that CHD4 enhances p65-mediated transcriptional activation of the Foxp3 promoter containing the CNS1/CNS2 regulatory regions. (e) Bisulfite sequencing analysis of the Foxp3 CNS2 enhancer (TSDR). Representative methylation patterns are shown for WT and CHD4-deficient Teff and Treg cells. Filled circles represent methylated CpG sites and open circles represent unmethylated CpG sites. (f) Quantification of Foxp3 CNS2 methylation. CHD4 deficiency induced a marked increase in DNA methylation within the TSDR specifically in Treg cells, whereas the methylation status of conventional T cells remained unchanged. (g-i) The survival of YFP+ CHD4-/- Tregs (1x10^6^) at 4 weeks post-adoptive transfer of Teffs (0.25x10^6^) was significantly decreased compared to the survival of WT Treg (*p<0.05, ***p<0.001), as shown by flow cytometric evaluation of viable cells. (i) Rag1-/- mice (5/group) received BALB/c cardiac allografts and adoptive transfer of WT Teff cells plus Tregs (2:1 ratio) from CHD4-/- or WT mice (Kaplan-Meier, *p<0.05). Data are presented as mean ± SD. Statistical significance was determined using two-tailed Student’s t-test. P values are indicated in the figure. LN, lymph node; SP, spleen; Teff, conventional CD4⁺ T cells; TSDR, Treg-specific demethylated region.

Since Treg lineage stability critically depends on the demethylated state of the Foxp3 CNS2 enhancer, we next assessed DNA methylation by bisulfite sequencing. Compared with WT Tregs, CHD4-deficient Tregs exhibited marked hypermethylation of the CNS2 CpG island, whereas no differences were observed in conventional T cells (Fig. 3e, f). Importantly, our group previously demonstrated that MBD2, a core component of the NuRD complex, recruits TET2 and TET3 to the Foxp3 CNS2 enhancer, thereby promoting CNS2 demethylation and stable Foxp3 expression. Together, these findings suggest that CHD4 is required to preserve the epigenetic landscape of the Foxp3 locus and raise the possibility that disruption of CHD4 compromises the ability of the NuRD complex to support MBD2/TET-dependent maintenance of the demethylated CNS2 enhancer, ultimately impairing stable Foxp3 expression and Treg identity.

To assess the effects of CHD4 deletion on Treg viability and function, we first tested the ability of Tregs to undergo homeostatic proliferation over 30 days post-adoptive transfer into immunodeficient mice. We found that the total numbers of yellow fluorescent protein–positive (CD4^+^YFP^+^) Tregs (Fig. 3g) and CD4+Foxp3+ Tregs (Fig. 3h, i) were markedly decreased in CHD4^-/-^ vs. WT Tregs, suggesting CHD4 deletion affected Treg survival. In a second *in vivo* test of CHD4 deletion in Tregs, we undertook cardiac allografts using BALB/c cardiac donors and immunodeficient B6/Rag1^-/-^ recipients (Fig. 3j). Immunodeficient recipients adoptively transferred with WT Teff and CHD4^-/-^ Tregs developed acute rejection by 14 d post-transplant, whereas mice receiving co-transfer of WT Teff and WT Treg cells maintained their allografts long-term (>100 d), further illustrating Treg impairment upon CHD4 deletion. These findings provide further evidence that CHD4 deficiency results in the loss of Foxp3⁺ Treg lineage identity *in vivo*.

### Rescue of CHD4^-/-^ mice by adoptive transfer of WT Treg cells

Next, we assessed the effects of adoptive transfer of WT Tregs to CHD4^-/-^ male mice that otherwise died by 3-4 weeks of life. Transfer of normal Tregs extended life span to >110 days (Fig. 4a, video 1), restored normal weight gain (Fig. 4b), and normalized IgG (Fig. 4c) and CD4, CD8, CD25, YFP, Foxp3 and Ki67+ levels (Fig. 4d, Fig. S3a). Interim analysis at 54 days post-transfer of circulating Tregs (CD4+CD25+YFP- cells, 0.26% and CD4+CD25+YFP+ cells, 6.68%) showed that ∼90% of the Tregs in the treated CHD4-/- mice were of exogenous origin (CD4+CD25+YFP- cells) (Fig. 4d). Screening of sera for autoantibodies showed that exogenous Tregs were able to control part of the autoreactive B cell clones (Suppl. Table 2). After euthanizing the adoptively transferred mice at day 110, we found that Foxp3 expression was decreased compared to WT mice, but compared to CHD4^-/-^ mice (Fig.1c), the treated group still showed an increase in Tregs in lymph nodes and spleen, with about half of the Tregs still coming from the originally injected WT Tregs (CD4+CD25+YFP- cells) (Fig. 4e-f). Compared to littermates, CHD4^-/-^ mice treated with WT Tregs showed increased CD4+CD44+CD62L^lo^ cells and CD4+ Ki67+ cells in lymph nodes and spleen (Fig. S3b, c), and modest impairment of *in vitro* suppressive function (P<0.05) (Fig. 4g). Adoptive transfer of WT Tregs also inhibited the production of most, but not all, autoantibodies detected in CHD4^-/-^ mice (Fig. S2, Suppl. Table 2). Hence, adoptive transfer of WT Tregs can overcome the exuberant T and B cell responses in CHD4^-/-^ mice for prolonged periods. These data are consistent with evidence that adoptively transferred Tregs are capable of achieving long-term persistence and stable expansion *in vivo*, and support the feasibility of Treg-based therapies in allergy, autoimmunity and transplantation (22).

**Fig. 4.**
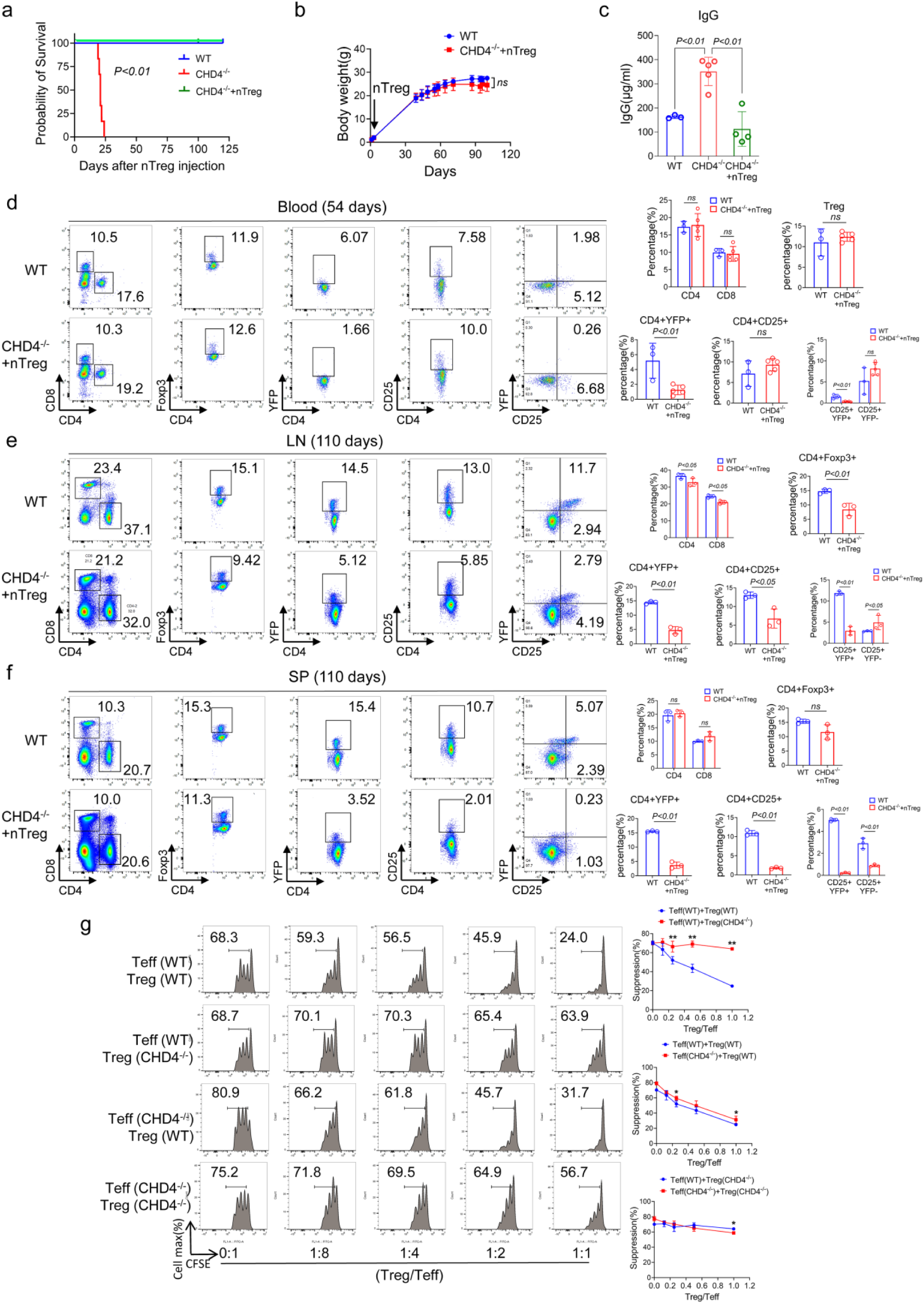
Recovery of CHD4-/- mice by adoptive transfer of WT Tregs (2.5x106, i.v.) at day 3 of life. (a) Survival and (b) body weight (n=5/group). (c) Serum IgG and (d) percentages of YFP+ cells and CD4+ cells at 54 d post-injection of WT Tregs (n=5/group). Percentages of YFP+ cells and CD4+ cells in (e) lymph nodes and (f) spleen at 100 d post-Treg injection vs. WT control (n=5/group). (g) Treg suppression assay using pooled Tregs from lymph nodes and spleens of WT and CHD4^-/-^ mice reconstituted with WT Tregs, with representative data shown on the right, along with the percentage of proliferating cells in each panel. Data are shown as mean ± SD, 5 samples/group. ANOVA; *p<0.05, **p<0.01 or ns (not significant) vs. control.

### Conserved CHD4 interactomes in Treg and conventional T cells

To determine whether the unique dependence of Treg cells on CHD4 reflected the assembly of lineage-specific protein complexes, we performed mass spectrometry analysis of CHD4 co-immunoprecipitates from Treg and conventional CD4⁺ T cells. Proteomic profiling revealed a substantial overlap in CHD4-interacting proteins between the two cell populations (Fig. S4a). Notably, a significant proportion of the identified interactors corresponded to canonical components of the NuRD complex, underscoring the central role of CHD4 as a core scaffold and functional hub within this chromatin-remodeling complex (Fig. S4b). Consistent with this, the NuRD and Sin3/HDAC complexes represented the predominant CHD4 interactomes in both Treg and Teff cells (Fig. S4c), with only modest differences in the relative abundance of individual interactors detected between the two populations. These findings indicate that the selective requirement for CHD4 in Treg biology is unlikely to result from the formation of Treg-specific CHD4-containing protein complexes. Rather, they suggest that the functional specificity of CHD4 is dictated by the distinct epigenetic landscape in which these conserved chromatin-remodeling complexes operate, thereby conferring a unique dependence of Treg cells on CHD4 activity. Collectively, these observations support CHD4 as an attractive and potentially druggable target for selectively modulating Treg function.

### Characterization of novel CHD4 inhibitors

CHD4 is a giant chromatin remodeling protein (250-280 kDa) with multiple domains, including helicase and adjacent tandem chromodomains, PHD domains, and ∼1000 N- and C-terminal residues encoding for unknown structures and functions (Fig. 5a) (23). With a translational perspective, we undertook cell target engagement studies using the Micro-Tag cell target engagement platform to discover novel CHD4 inhibitors (24, 25). This platform measures transitions in the thermodynamic state of target protein during engagement with drug molecules (26), and relies on split-RNAse A that enables interrogation of drug targets without interfering with their folding, localization or function (24, 25). Using MICRO-TAG reporter cells optimized for CHD4, a workflow was designed for screening, identification and characterization of CHD4 inhibitor candidates (Fig. 5b). We identified the thermal melting profile of CHD4 and found the half-way temperature of aggregation (Tagg50) to be 50 ℃ (Fig. 5c). The temperature point of 37 ℃ represented the physiological temperature at which CHD4 would exist in its native structural conformation with all inherent protein-protein interactions, whereas the temperature point of 50℃ represented Tagg50 at which CHD4 would exist in half-unwound structural conformation, thereby providing unblocked access to its ligandable hotspots. Therefore, nomination of the top drug CHD4 inhibitor candidates was based on their enrichment at 35 ℃ vs 50 ℃ selections. Candidates CH 41, CH 61 and CH 81 showed target-stabilizing profiles at 50 ℃ selection as compared to the 35 ℃ selection. All other candidates that did not follow this rationale were disqualified. The closely related compounds CH41, CH61 and CH81 were selected for further characterization. All three candidates were tested at 2.5 µM using real-time MICRO-TAG cellular target engagement in the temperature series format, and each demonstrated direct engagement with CHD4 (Fig. 5d). Their target engagement profiles were interpreted as stabilization of CHD4, as judged by increased signal at temperatures above 50 °C, consistent with the above determined Tagg50 for CHD4. The data demonstrate that CH41 and its analogs are cell permeable with ability to directly engage CHD4. Deeper dose-dependent profiling of the three candidates using MICRO-TAG real-time cell target engagement led to nomination of CH41 as our top CHD4i. CH41 demonstrated dose-dependent cell target engagement with EC50 of 761 nM with full-length CHD4 (Fig. 5e). To gain further proof of target engagement, we measured CHD4 abundance in response to CH41 treatment. In comparison to controls, CH41 reduced the abundance of CHD4 by 1.2x10^7^ fluorescence units at the 24 h time point, suggesting cellular recycling of CHD4 that is inhibited and conformationally stabilized by CH41 (Fig. 5f). Together, these data establish that CH41 robustly and reproducibly engages CHD4 in cells, providing a validated chemical framework for investigating the role of CHD4 and NuRD modulation (23, 27).

**Figure 5.**
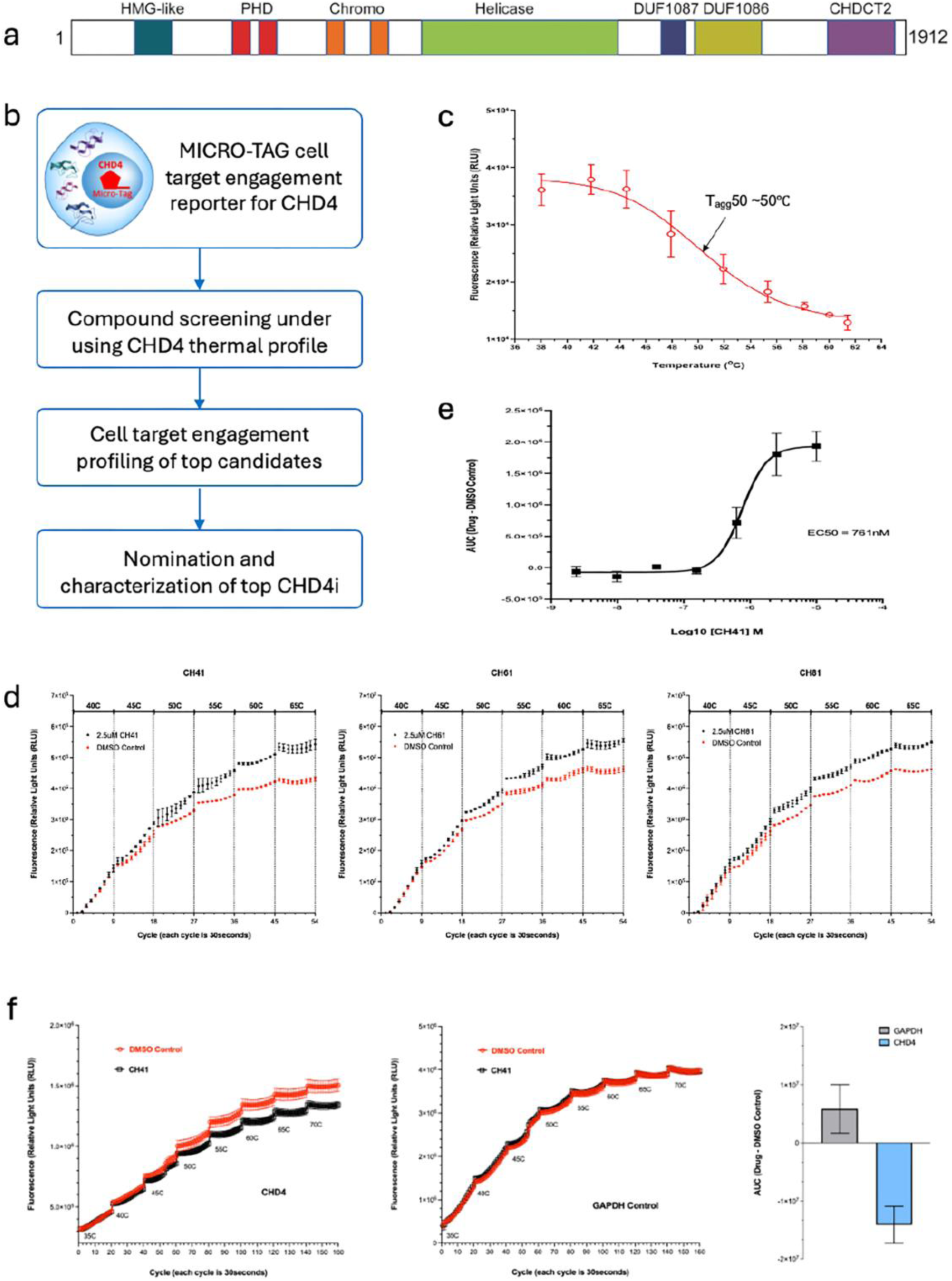
Identification and characterization of CHD4 inhibitors. (a) Domain architecture of full-length CHD4, illustrating its multi-domain organization. (b) MICRO-TAG workflow using engineered CHD4 reporter cells for real-time cellular target engagement. Cell lines expressing full-length CHD4 were generated to enable quantitative measurement of CHD4 engagement in cells. **(c)** Thermal melting profile of full-length CHD4 in intact cells, quantified using the MICRO-TAG cellular target engagement assay. The midpoint of stabilization (Tagg50) reflects the point under which CHD4 exists 50% structurally destabilized state. (d) Real-time MICRO-TAG cellular target engagement of CHD4 by the top three CHD4i candidates, CH41, CH61 and CH81, as measured at a single dose and multiple temperature points. The compounds exhibit comparable engagement profiles, indicating that CHD4 stabilization is a class property of this probe series. (e) Potency of cellular target engagement (EC50) of CHD4 with CH41, the selected to CHD4i, as measured employing real-time MICRO-TAG cellular target engagement. **(f)** Real-time cell target engagement data showing reduced abundance of CHD4 induced by CH41 treatment within 24 hours.

### CHD4 inhibitor impairs murine Treg function phenocopying Chd4 knock-out

Subsequent *in vitro* studies showed that CHD4i (CH41) decreased Treg expression of Foxp3 in Tregs (Fig. 6a), led to lower levels of ac- H3K27 binding at the Foxp3 promoter compared to DMSO treatment (Fig. 6b), and impaired expression of other Treg-linked genes such *Cxcl4, Ccr7, Il10 and Ikzf2* (Fig. 6c). Treatment with CHD4i also reduced TGF-β-induced conversion of naïve CD4⁺CD25⁻ T cells into Foxp3⁺ regulatory T cells (Fig. 6d) and impaired the suppressive function of Treg cells (Fig. 6e). Notably, CHD4 inhibition with CH41 did not disrupt the assembly of the NuRD complex, indicating that the observed functional effects result from inhibition of CHD4 activity rather than destabilization of the complex (Fig. 6f).

**Fig. 6.**
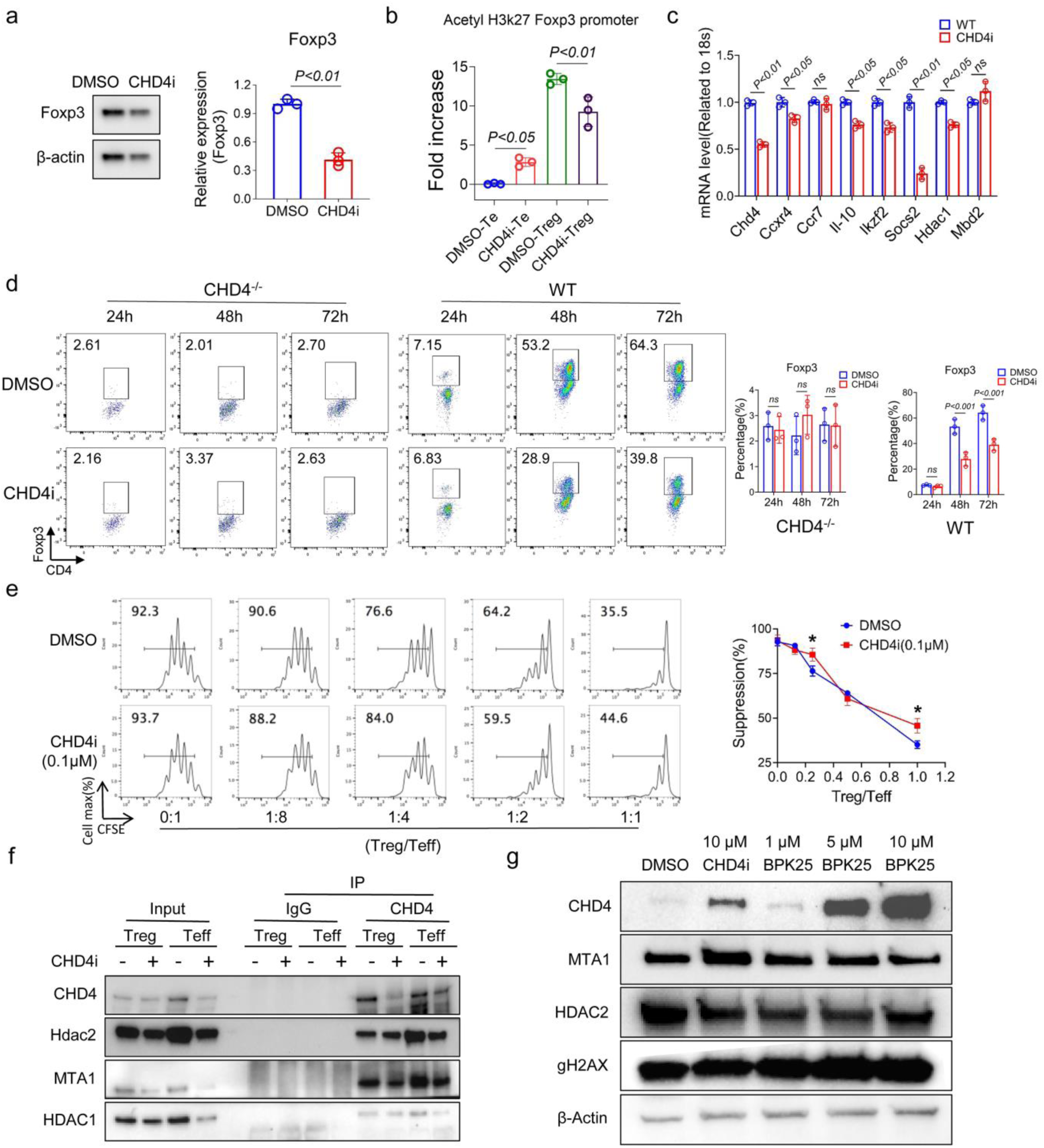
Effects of CHD4 inhibitor, CHD4i, on Treg gene expression and function *in vitro*. (a) Representative bands (left) and statistical analysis (right) of Western blotting for Foxp3 expression in freshly isolated Tregs treated with CHD4i or DMSO. (b) ChIP-qPCR for H3K27ac binding to Foxp3 promoter. (c) qPCR analyses of indicated gene expression in T effector and Treg cells. qPCR data were normalized to 18S (in (a-c) *p<0.05, **p<0.01, ***p<0.001, ****P < 0.0001 vs. WT control), and data (mean ±SD) are representative of 2 independent experiments. (d) Graphs report the iTreg conversion analysis after the CHD4i treatment for 24 h, 48 h and 72 h of Teff isolated from spleen and lymphocyte of Foxp3YFP/Cre mice (on the left) or CHD4^-/-^ mice (on the right). Histograms of the average of two different experiments of iTreg conversion on Teff from control cells and one on Teff derived from CHD4 -/- mice. (e) Treg suppression assay using pooled Tregs from lymph nodes and spleens of WT and CHD4^-/-^ mice, with representative data shown in the right, along with the percentage of proliferating cells in each panel. (f) Western blot representing the co-IP of Tregs and Teff after the treatment with CHD4i/DMSO. IP was performed for CHD4. Indicated proteins were developed. IgG was used as negative control of IP.

### Genetic and chemical inhibition of CHD4 induce concordant histone post-translational modification (PTM) redistribution

To define the role of CHD4 in regulating histone PTM landscapes and to assess effects of a CHD4 inhibitor, we compared the effects of genetic and chemical CHD4 perturbation on histone PTM composition. Prior studies have demonstrated that genetic depletion of CHD4 or other NuRD components leads to altered histone PTM landscapes, including increased histone acetylation and redistribution of methylation states at regulatory loci, consistent with loss of NuRD-associated HDAC activity and remodeling function (28–30). Genetic depletion of CHD4 was used to establish the chromatin modification changes associated with loss of CHD4 function, while treatment with the small-molecule inhibitor CH41 was evaluated to determine whether chemical inhibition phenocopies these effects. By directly comparing histone PTM redistribution following sgCHD4-mediated knockdown and CH41 treatment, this approach enabled assessment of mechanistic concordance and provided a functional benchmark for determining whether CH41 phenocopies CHD4 genetic loss at the level of chromatin regulation. Comparative analysis of histone PTMs revealed that genetic depletion of CHD4 (sgCHD4) and chemical inhibition with CH41 produced broadly similar patterns of PTM redistribution across multiple histone residues (Fig. S5). For the majority of marks, fold changes clustered nearly identically, indicating that neither perturbation globally disrupts histone modification levels but instead induces selective redistribution among modification states. Notably, both sgCHD4 and CH41 treatment elicited concordant directional changes at several residues associated with chromatin regulation, including shifts in lysine acetylation and methylation states on histones H3 and H4. While chemical inhibition with CH41 generally produced larger-magnitude effects than genetic depletion, the overall pattern of affected PTMs was highly overlapping, consistent with on-target inhibition of CHD4-dependent chromatin regulatory activity. Larger fold changes observed for low-abundance acetylation marks (*e.g.* H1.4: K25, H2A: K36, H2A3: K15) likely reflect amplification effects inherent to relative abundance measurements and are therefore interpreted as directional rather than absolute changes. Collectively, these data support a shared chromatin regulatory signature between genetic and chemical CHD4 inhibition, consistent with CH41 phenocopying key aspects of CHD4 loss at the level of histone PTM organization.

### CHD4 inhibitor promotes antitumor immunity

Given data suggesting that CHD4i preferentially affected the functions of Tregs versus Teff cells (Fig. 6), and TCGA analysis showing increased CHD4 expression and FOXP3+ Tregs within lung tumors (Fig. S6), we considered the application of CHD4i in cancer. We used two syngeneic cancer models to assess the effects of CHD4i on Treg function *in vivo*. First, C57BL/6 mice injected with TC1 lung adenocarcinoma cells were randomized into groups and injected daily with CHD4i for ten days at 2.5 mg/kg or 5 mg/kg. CHD4i treatment with 5 mg/kg significantly decreased TC1 tumor growth in immunocompetent (Fig. 7a, b) but not immunodeficient (Fig. 7c, d) C57BL/6 mice. CHD4i therapy decreased intratumoral Foxp3 Treg cells (Fig. 7e), increased intratumoral IL-2 production (Fig. 7f), and decreased CD4+ and CD8+/PD1+ cells at the tumor site and within spleens of mice bearing TC1 tumors (Fig. 7g). These data demonstrated the ability of CHD4i to reactivate anti-cancer adaptive immunity. We then assessed the effects of CHD4i in a second tumor model involving orthotopic HCC tumors established in BALB/c and Rag1-/- mice by intrahepatic injection of H22 cells (Fig. 8). CHD4i treatment (5 mg/kg/d) significantly inhibited HCC proliferation in immunocompetent (p<0.05) (Fig. 8a, b) but not immunodeficient mice (Fig. 8c, d), with a decrease in AFP levels in sera of WT mice tumor-bearing treated with CHD4i (Fig. 8e). The intratumoral percentage of Foxp3 Treg cells was decreased in the CHD4i group compared with DMSO (Fig. 8f) and without changes in CD4 and CD8 T cells (Fig. S7a). However, tumor-infiltrating Ki-67+ CD4 T cells increased from 42.2% to 58.1% (n=5, p<0.05) (Fig. S7b), along with increased IL-2 expression in tumor-infiltrating T cells (CD4, n=5, p<0.01 and CD8, n=5, p<0.05) (Fig. 8g), with no differences in spleen or lymph node T cells (n=5) (Fig. 8h). In addition to IL-2, IFN-γ expression was increased in tumor-infiltrating T cells (CD4, n=5, p<0.05 and CD8, n=5, p<0.05) and in Tregs increased from 22.18% to 38.2% (n=5, p<0.05) (Fig. 8h).

**Fig. 7.**
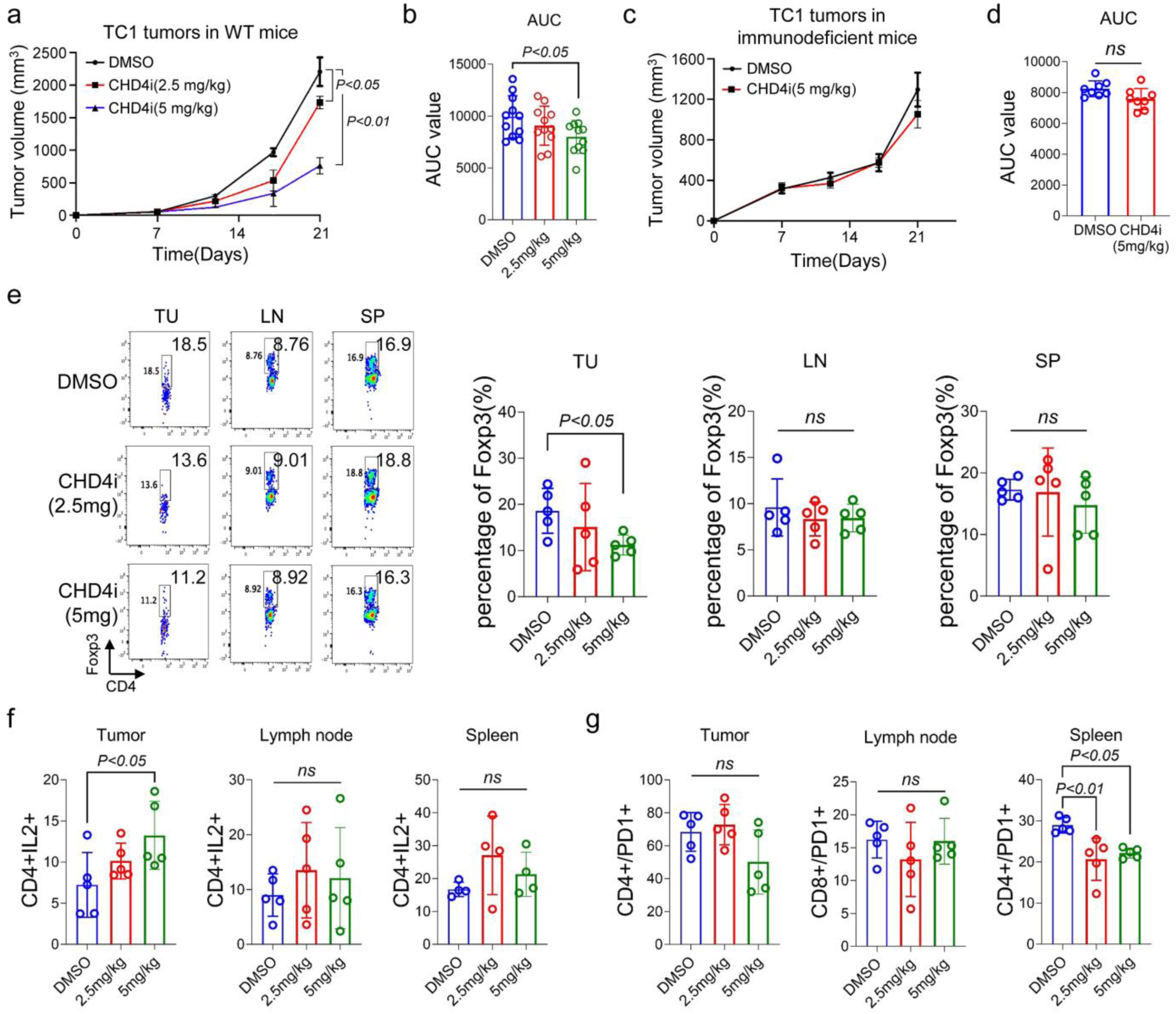
CHD4i enhances anti-tumor immunity (lung tumor model). Decreased (a) TC1 tumor volumes and (b) area- under-curve data in C57BL/6 mice treated with CHD4i (2.5 or 5 mg/kg/day) vs. DMSO (n=8-10/group) (*p<0.05), whereas (c) TC1 tumor volumes and (d) area-under-curve data were comparable in Rag1^-/-^ mice treated with CHD4i (5 mg/kg/day) vs. DMSO (n=8-10/group) (*p<0.05). Percentages of (e) CD4+Foxp3+ cells, (f) CD4+IL2+ cells, CD4+IL2+ cells and (g) PD-L1+CD4+ cells, PD-L1+CD8+ cells in lymph nodes, spleens and tumors from CHD4i and DMSO-treated groups. Data are shown as mean ± SD, 8-10 samples/group. ANOVA; *p<0.05, **p<0.01 or ns (not significant) vs. control.

**Fig. 8.**
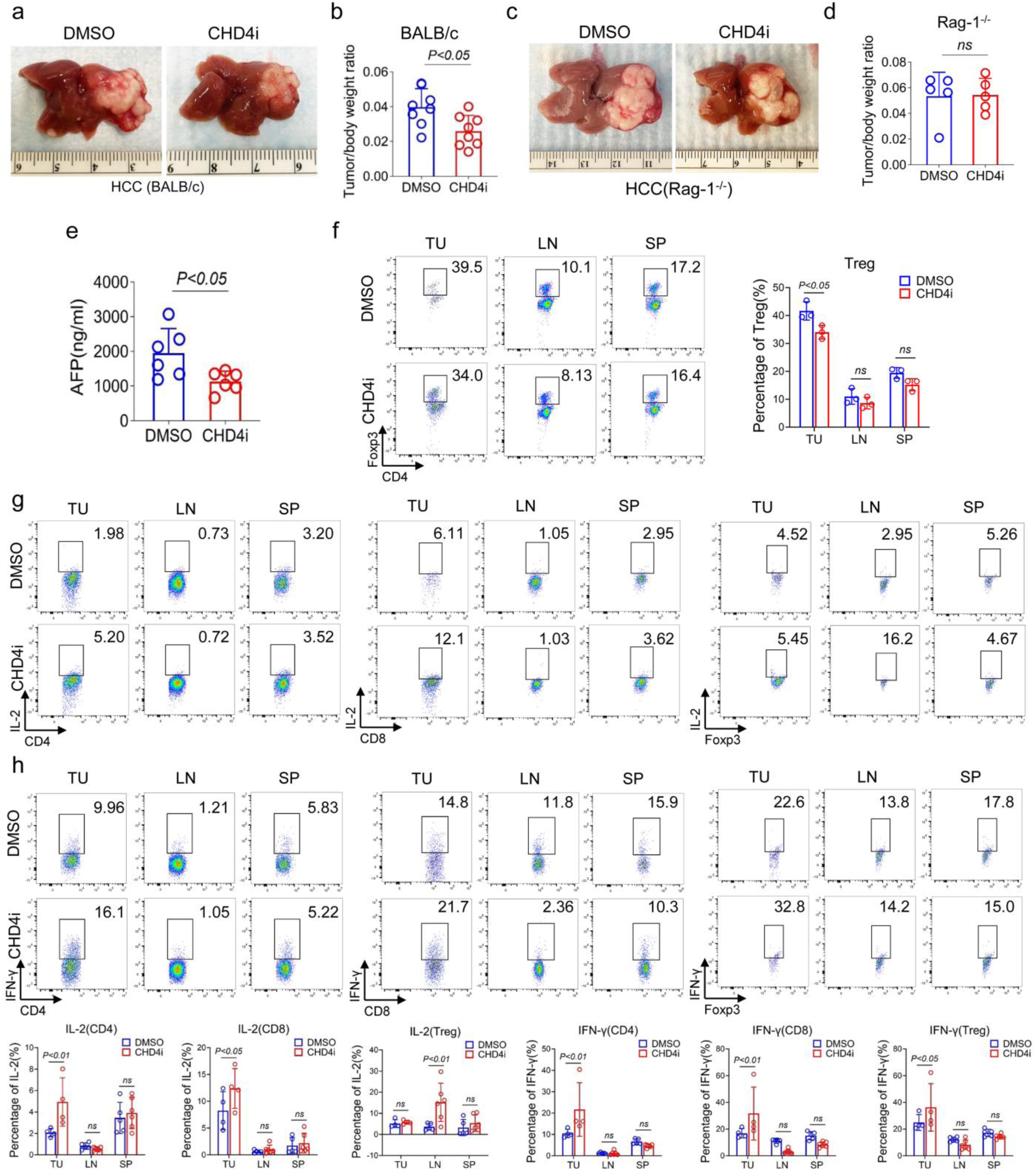
CHD4i enhances anti-tumor immunity in a syngeneic orthotopic HCC tumor model. Decreased (a) tumor volumes and (b) weights in mice treated with CHD4i (5 mg/kg/day) vs. DMSO (n=5-8/group) (*p<0.05). (c) Tumor volumes and (d) weights were comparable in Rag1-/- mice treated with CHD4i (5 mg/kg/day) vs. DMSO (n=5/group). (e) Serum AFP levels of CHD4i vs. DMSO-treated groups. Percentages of (f) CD4+Foxp3+cells, (g) CD4+IL2+ cells, CD8+IL2+ cells, Foxp3+IL2+ cells and (h) CD4+IFN-γ+ cells, CD8+IFN-γ+ cells, Foxp3+IFN-γ+ cells in lymph nodes, spleens and tumors of CHD4i vs. DMSO-treated groups. Data are shown as mean ± SD, 8-10 samples/group. Student’s t- test for unpaired data; *p<0.05, **p<0.01 or ns (not significant) vs. control.

Hence, in a second tumor model, CHD4i promoted T cell infiltration and cytokine production while decreasing Treg cell infiltration and promoting Treg dysfunction as indicated by their increased IL-2 and IFN-γ expression. Of note, these studies showed modulation of Treg and Teff cell responses within tumors but not secondary lymphoid tissues, and were not associated with cell infiltrates in solid organs (liver, lung, kidney, skin histology), or biochemical abnormalities of renal (BUN, Cr) or liver (AST, ALT) function.

## DISCUSSION

The activation and functions of Treg cells are controlled through various pathways, including the actions of nuclear protein complexes that govern immune responses via epigenetic mechanisms (31). CHD4, a core component of the NuRD complex, is known to play an essential part in lymphocyte generation and differentiation (14, 16, 17). In the current study, we identified CHD4 as a critical regulator of Foxp3 expression and Treg functionality. We found that CHD4 bound to Foxp3 and Tip60, but not p300 (Fig. 1a), and have previously shown that IP of Foxp3 led to co-IP of Tip60 in Tregs, which is important for Foxp3 acetylation, dimerization, and Treg function (21). Hence, we undertook conditional deletion of CHD4 in Treg cells. Such CHD4 deletion resulted in lethal autoimmunity, characterized by severe anemia, splenomegaly and lymphadenopathy, mononuclear cell infiltrates, and auto-antibody production, similar to the phenotype of mice with mutations of Foxp3 (32, 33). These data, plus our findings using a newly developed CHD4 inhibitor in WT mice, underscore the potential benefit of targeting this protein for therapeutic purposes, such as in cancer immunotherapy.

FOXP3 is the master control gene for Treg development and function and is regulated by histone acetylation and CNS2 methylation (34). We found CHD4 deletion decreased Tregs in peripheral lymphoid tissues, reduced Foxp3 expression in Tregs, allowing increased T-cell activation and production of proinflammatory cytokines (Fig. 1). Our ChIP studies show CHD4 bound to the CNS1 in Treg cells (Fig. 3) and CHD4 KO Tregs showed a highly methylated phenotype, with methylation of almost all of the CNS2 CpG sites (Fig. 4). Likewise, CHD4 targeting Tregs decreased TGF-β-induced conversion of naïve CD4^+^CD25^−^ T-cells into Foxp3^+^ iTregs, Hence, CHD4 deletion altered the epigenetic regulation of Foxp3 expression and impaired Treg cell function.

As a key member of the NuRD complex, CHD4 can interact and cooperate with other NuRD components, including Mbd2, Hdac1 and Hdac2, but can execute its functions alone or as part of additional high-molecular weight complexes (15, 35). Our RNA-seq analysis comparing CHD4-, Hdac1-, and Hdac2-deficient Tregs demonstrated that the transcriptional consequences of CHD4 deletion closely resemble those observed following Hdac2 deletion, particularly when analysis was restricted to the canonical Treg gene signature. These findings support the concept that CHD4 exerts its functions in Treg cells primarily through the NuRD complex. To determine whether the selective dependence of Tregs on CHD4 could instead reflect the assembly of lineage-specific CHD4-containing protein complexes, we performed mass spectrometry analysis of CHD4-associated proteins in Tregs and conventional CD4⁺ T cells. Surprisingly, CHD4 displayed a highly conserved interactome in both cell populations, with the NuRD and Sin3/HDAC complexes representing the predominant CHD4- associated complexes and only minor differences in the abundance of individual interactors. Thus, the unique vulnerability of Tregs to CHD4 depletion is unlikely to result from the formation of Treg-specific CHD4 protein complexes. Rather, our findings suggest that Treg dependency arises from the distinct epigenetic programs controlled by CHD4-containing chromatin-remodeling complexes, to which the Treg lineage is particularly dependent for the maintenance of FOXP3 expression, lineage stability and suppressive function.

These observations identify CHD4 as a key epigenetic node whose pharmacological inhibition selectively disrupts the Treg transcriptional program despite its association with conserved chromatin-remodeling complexes in conventional T cells. We identified CH41, a novel and potent inhibitor of CHD4, using our cellular target engagement platform (Fig. 8). We first focused on testing effects of CHD4i on Tregs and conventional T cells *in vitro* (Fig. 9). We found that CH41 decreased expression of several signature genes in Foxp3+ Tregs, including genes encoding CTLA4, Foxp3, GITR and TGF-β, increased IFN-γ and IL-17 expression, and drastically reduced the conversion of Teff cells to Foxp3+ iTreg cells, highlighting an important role of CHD4 in Treg development and functionality. Subsequent use of CH41 in tumor models led to significant inhibition of tumor growth without evidence of concomitant development of autoimmunity. We conclude that while tumor-associated Tregs have unique properties that include their increased suppressive activity compared to non-tumor associated Treg cells (11, 36), they are also more susceptible than the latter and conventional T cells to pharmacologic targeting of the CHD4 chromodomain. Such findings provide a rationale for further drug development around CHD4 in the context of cancer immunotherapy.

In summary, our data demonstrate that loss of CHD4 in regulatory T cells results in increased DNA methylation at the Foxp3 CNS2 region, a key epigenetic hallmark associated with impaired Foxp3 stability. Importantly, this alteration occurs in the context of widespread redistribution of histone PTM induced by CHD4 deficiency. Given the established crosstalk between histone modifications and DNA methylation, we propose that disruption of CHD4-dependent chromatin organization leads to an altered histone modification landscape at the Foxp3 locus, thereby permitting aberrant DNA methylation at CNS2. Such epigenetic reprogramming likely compromises the maintenance of Foxp3 expression, ultimately resulting in reduced Foxp3 protein levels and loss of Treg identity.

In a broader context, our work also provides insights into why Foxp3+ Tregs have three nuclear complexes, NuRD, Sin3 and CoREST, that each contain Hdac1 and/or Hdac2 that are catalytically active only when embedded in these large coregulatory complexes (31). Each complex brings distinct scaffolds, readers, and interactors that let the same catalytic core (Hdac1/2) execute different chromatin programs at different loci, times, and activation states. In Tregs, NuRD (current data), SIN3 (37), and CoREST (38) are not redundant; they control partly non-overlapping aspects of Foxp3 expression, Treg lineage stability, and repression of effector genes. Ultimately, the NuRD complex is key to Treg development, SIN3 is central to maintenance of Treg stability, and CoREST helps regulate Treg functions in the context of inflammatory responses.

## METHODS

### Sex as a biological variable

Our study examined male and female animals, and similar findings are reported for both sexes.

### Mice

We used male WT BALB/c, WT C57BL/6, Rag1^−/−^ C57BL/6, Rag1^−/−^ BALB/c and CD90.1/B6 mice from The Jackson Laboratory, plus previously described Foxp3*^YFP-cre^* mice (39) and CHD4*^flox/flox^* mice (40). Mice were housed under specific pathogen-free conditions, with 12-h light/dark cycles and access to food and tap water and studied with protocols approved by the Institutional Animal Care and Use Committee of the Children’s Hospital of Philadelphia (CHOP).

### Complete blood count

Blood was drawn from WT C57BL/6 and CHD4-/- mice and sent for complete blood count (CBC) analysis at the CHOP Hematology Core Lab.

### Plasmids and chemical compounds

We purchased the plasmids Flag-CHD4, Flag-Foxp3 and HA-p300 from Addgene for transient expression in 293T cells. CHD4i compound (CH41) was synthesized by E.N. but its chemical characterization is currently undergoing patent review prior to disclosure.

### Detection of autoantibodies

We analyzed sera of four CHD4-/- male mice at 21 days of life and four FOXP3-Cre control male mice at the same age. Additionally, sera of five Treg-treated mice were analyzed at days 42, 54 and day 110. Sera were diluted 1/10 with PBS and incubated with 4-5 µm cryosections of corresponding tissues from B6 WT mice for 1 hour, then washed twice in PBS. After washing, slides were incubated for 45 min with Alexa Fluor® 488 AffiniPure F(ab’)₂ Fragment Goat Anti-Mouse IgG (H+L), catalog #115-546-062, Jackson ImmunoResearch Lab, washed x2 in PBS and mounted with SlowFade Diamond Antifade Mountant (Invitrogen, #S36967). In the case of pancreas, sera were used in 1/1 dilution, as required to evaluate a presence of autoantibodies to islet cells. Mouse sera with known patterns of autoantibodies served as positive controls, while secondary anti-mouse antibody alone, WT sera and sera from FOXP3-Cre control mice served as negative controls.

### MICRO-TAG reporter cell line for CHD4

Micro-Tag reporter cells for CHD4 were generated by cloning the target gene sequence (UNIPROT ID: Q14839) in-frame with the sequence KETAAAKFERQHMDS at the C-terminus in pcDNA3 plasmid. This tag corresponds to the S-tag peptide derived from bovine pancreatic ribonuclease A (RNase A). The S-tag is the small subunit of a two-subunit ribonuclease S (RNase S). RNase consists of the two proteolytic fragments of bovine pancreatic ribonuclease A (RNase-A): the S-peptide (residues 1-20) and S-protein (residues 21-124). The Micro-Tag cell target engagement system was assembled in accordance with recent reports (24, 25, 41).

### Lysis preparation for MICRO-TAG enzyme complementation

HEK293 cells transiently expressing tagged CHD4 were grown in a 6-well tissue culture plate. The cells were lifted by pipetting up/down and transferred to a 15 ml tube. Cells were pelleted at 400 g for 3 min, washed with TBS (Tris Buffered Saline (1X TBS; 150 mM NaCl, 50 mM Tris-HCl pH 7.4) (Thermo Fisher Scientific; cat J62938.k7) and pelleted again. TBS was removed and cells lysed with 300 ul non-denaturing lysis buffer (non-denaturing Lysis Buffer: 1% Triton X-100 (in TBS) containing cOmplete, Mini, EDTA-free Protease Inhibitor Cocktail (Roche; #11836153001) for 1 h, at 4 °C on a rotator. The lysate was clarified by centrifugation at 14,000 rpm for 1 min in a microcentrifuge.

### Immunoblot analysis for MICRO-TAG CHD4

Immunoblot analysis was carried out using standard procedures. HEK293 cells transiently expressing tagged CHD4 of interest after 48 h were transferred to a 15 ml tube, pelleted at 400 g for 3 min, washed with TBS (Tris Buffered Saline (1X TBS; 150 mM NaCl, 50 mM Tris-HCl pH 7.4) (Thermo Fisher Scientific; cat J62938.k7) and pelleted again. TBS wash was removed, and cells lysed with 300 ul non-denaturing lysis buffer consisting of 1% Triton X-100 (in TBS) containing cOmplete, Mini, EDTA-free Protease Inhibitor Cocktail (Roche; cat #11836153001) for 1 h at 4 °C on a rotator. Lysates were clarified by centrifugation at 14,000 rpm for 1 min in a micro-centrifuge and used for Western analysis with S- Tag (D2K2V) XP rabbit mAb #12274.

### MICRO-TAG temperature series cell target engagement assay for CHD4 employing non-denaturing lysates

Test ligands were prepared in non-denaturing lysates were diluted 1/20 with cold TBS on ice and aliquoted 49.5 ul/well to 12 wells of a 96-well PCR plate and 0.5 ul of the 100X diluted ligands was added. The 60 ul of reaction buffer containing S-protein and FRET RNA substrate were added to a MicroAMP Optical 96-well reaction plate and 20 ul of lysates + ligand added and FAM fluorescence (excitation 493 nm; emission 517 nm) detected with the temperature series programming (41).

### Enzyme complementation analysis

Non-denaturing lysates (20 ul) prepared from cells expressing S-Tag constructs were used with reaction buffer (60 ul). The 60 ul of reaction buffer containing S-protein and FRET RNA substrate were added to a MicroAMP Optical 96-well reaction plate and 20 ul of lysates added and FAM fluorescence (excitation 493 nm; emission 517 nm) detected for 30 min.

***Quantification and statistical analysis of CHD4 cell target engagement.*** Statistical analyses were performed using GraphPad Prism v.9 software. Data were analyzed using two-way ANOVA followed by a t-test. Statistical significance was considered when p< 0.05. All results are shown as mean ± SEM. Calculation of AUC (Area Under the Curve) was performed using GraphPad Prism software, and delta AUC determined by subtracting DMSO AUC from Drug treated AUC curve. Calculation of “observed in-cell dissociation constant KD” was by analysis of saturation binding using one-site total binding equation (One-site total binding model: Y=Bmax*X/(K_D_+X) + NS*X + Background) where background non-specific binding is constant set to zero, Y is the fluorescence relative light units (RLU), X is time (cycles: 30 sec each cycle), Bmax is the maximum specific binding in RLU, and NS is slope of non-specific binding (RLU/cycle).

### Treg suppression assays

For *in vitro* studies, 5×10^4^ cell-sorted CD4^+^CD25^-^ T cells and CD4^+^CD25^+^ Tregs from Foxp3^YFP-Cre^ and CHD4^-/-^mice isolated using CD4^+^CD25^+^Treg isolation kit (Miltenyi Biotec, #130-091- 041), or B6 Tregs using DMSO or CHD4i (0.1 μM) were added to 96-well plates. Equal numbers of CFSE-labeled CD4^+^CD25^−^ T cells and γ-irradiated antigen-presenting cells (APC), isolated using a CD90.1 kit (Miltenyi Biotec, #130-049-101), plus CD3 mAb (1 μg/ml), were cultured for 72 h. After 72 h, proliferation of Teff cells was determined by flow and analysis of CFSE dilution. For *in vivo* Treg suppression assays, 1×10^6^ CD4^+^CD25^−^ Thy1.1+ and 0.5×10^6^ Tregs were injected via the tail vein into Rag1^−/−^ mice. A week later, lymph node and spleens were harvested and stained with Thy1.1-PE and CD4-Pacific blue, and the numbers of Thy1.1^+^ T cell cells were determined using a CytoFlex flow cytometer.

### Flow cytometry and cell sorting

Single-cell suspensions from lymph nodes, spleens or tumors were prepared as described(38) and stained with fluorochrome-conjugated mAbs (BD Biosciences unless specified) that were directed against CD4 (Pacific blue, Invitrogen, #MHCD0428), CD8 (Super Bright 645, eBioscience, clone 53-6.7,#64-0081-82), FOXP3 (eFluor 450, eBioscience, clone FJK-16s, #48-5773-82), CD62L (PE-Cy7, clone MEL-14, #25-0621-82), IFN-γ (APC, clone XMG1.2, #554413), CD44 (PE-Cyaine5, eBioscience, clone IM7, #15-0441-83), and CD25 (APC, eBioscience, clone PC61.5, #17-0251-82) and acquired with a CytoFlex flow cytometer (Beckman). We purchased unconjugated CD3 (clone 145-2C11, #553057) and CD28 (clone 37.51, #553294) mAbs from BD. For cell sorting, we isolated lymphocytes from Foxp3YFP/Cre mice and purified CD4+ cells as above, followed by sorting CD4+YFP+(Foxp3+) and CD4+YFP− cells via a FACS Aria or FACS Aurora cell sorter (UPenn Cell Sorting Facility).

### iTreg conversion assay

Teff cells from Foxp3YFP/Cre mice or CHD4^-/-^ mice were isolated through the CD4+CD25+ regulatory Treg kit (Miltenyi Biotec) as described above. Teff cells were cultured from 24-72 h with a 1:1 ratio of CD3/CD28 mAb-coated beads, IL-2 (1 µg/ml) and TGF-β (3 ng/ml). Teff cells were also treated with CHD4 inhibitor (CHD4i) added in the medium (10 μM). Negative controls were treated with DMSO. After the required time, cells were collected and analyzed using a Cytoflex (Beckman Coulter) flow cytometer.

### Methylation study

DNA was purified (DNA kit; Qiagen) from isolated and sorted Tregs and Teff from Foxp3YFP/Cre mice or CHD4^-/-^ mice(34). After nested PCR for the Foxp3 CNS, bisulfite-conversion was performed (MethylDetector, Bisulfite Modification Kit, Active Motif), and the PCR product was cloned through TA cloning following the manufactures instructions (Quick PROTOCOL pGEM-T and pGEM-T Easy Vector Systems). Briefly nested PCR was performed in two steps with the following primers:

**FOXP3 CNS INNER FW**: 5’ TTGAGTTTTTGTTATTATAGTATTTGAAGAT
**FOXP3 CNS INNER RV**: 5’ ACTAAAAACCTAAAAAACTAAACTAACCAA
**FOXP3 CNS OUTER FW**: 5’ GGGTTTTGGGATATTAATATATATAGTAAG
**FOXP3 CNS OUTER RV**: 5’ CCACTATATTAACTTAACCCATATAACTAA

Bisulfite conversion was performed following the manufacturer’s instructions for 5 h; ligation using T4 DNA ligase was followed by transformation of JM109 competent cells and selection of colonies on LB/ampicillin/IPTG (0.5 mM)/X-Gal (80 µg/ml) plates. Colonies were grown and DNA was purified (Qiagen) and sequenced at the Penn Genomic and Sequencing Core (University of Pennsylvania).

### Microarray and real-time qPCR and GSEA analysis

RNA was isolated using RNeasy kits (QIAGEN), and RNA integrity and quantity were analyzed by NanoDrop ND-1000 and Nanochip 2100 Bioanalyzer (Agilent Technologies). Microarray experiments were performed using whole mouse genome oligoarrays (Mouse430a, Affymetrix) and array data analyzed using MAYDAY 2.12 software (42). Array data were subjected to robust multiarray average (RMA) normalization and analyzed using Student’s t test. Only data with a false discovery rate–adjusted P value of less than 0.05 and at least 2× differential expression were included in the analysis. Data underwent z-score transformation for display. The microarray data were deposited in the Gene Expression Omnibus (GEO) database (accession number GSE48653). Expression of individual genes was verified by qPCR. RNA was reverse transcribed to cDNA (Applied Biosystems) and qPCR performed using Taqman primer and probe sets; data were normalized to endogenous 18s rRNA, and relative expression was determined by the formula 2–ΔCT.

### Co-immunoprecipitation (Co-IP) and Western blotting

Treg and Teff cells from B6 mice were collected after treatment for 24 h at 10 µM with CHD4i and after centrifugation and washes with PBS 1X were lysed with RIPA Buffer (ThermoScientific) plus (PIC) 100 X (ThermoScientific) for 20 min at 4 °C on the rotor. Immunoprecipitation was performed with 4 µg of IgG rabbit and 4 µg of CHD4 antibodies (Cell Signaling D8B12). Samples were incubated overnight on the rotor at 4 °C. The next day magnetic bead-coupled protein G (Invitrogen) was added to each sample for 1 h and 30 min on the rotor at 4 °C. After washes with RIPA Buffer, magnetic beads were collected with magnetic support and lysed with Laemmli buffer 2X plus β-mercaptoethanol (Biorad) and PIC 100 X. After boiling, protein were separated by SDS-PAGE, transferred to polyvinylidene difluoride (PVDF) membranes (BioRad), and analyzed using antibodies against CHD4 (Cell Signaling D8B12, Cell Signaling D4B7), HDAC1 (Abcam, ab53091), HDAC2 (Cell Signaling), Mta1(Cell Signaling D17G10), Foxp3 (eBioscience, FJK-16s). Secondary HRP-conjugated Abs against mouse (7076), and rabbit (catalog 7074) IgG were purchased from Cell Signaling. Streptavidin HRP antibody was purchased from BD (554066).

### Mass spectroscopy

Treg and Teff cells from WT mice were isolated from spleen and LN and sorted as previously described. After centrifugation and washes with PBS 1X they were lysed with RIPA Buffer (ThermoScientific) plus (PIC) 100 X (ThermoScientific) for 20 min at 4°C on the rotor. Immunoprecipitation was performed with 4 µg of IgG rabbit and 4 µg of CHD4 antibodies (cell signaling D8B12 and cell signaling D4B7). Samples were incubated overnight (on) on the rotor at 4 °C. On the next day, magnetic bead-conjugated protein G (Invitrogen) was added to each sample for 1 h and 30 min on the rotor at 4 °C. After washes with Buffer A (20 mM Tris-HCl pH 7.5, 1.5 mM MgCl2, 150 mM NaCl), magnetic beads were collected with magnetic stand and 90% of beads were sent for Mass Spectrometry Analysis to the Proteomic Core at the University of Pennsylvania and subjected to Orbitrap preparation procedure. The remaining 10% was lysed with Laemmli buffer 2X plus β-mercaptoethanol (BioRad) and PIC (100 X); after boiling, proteins were separated by SDS-PAGE and transferred to polyvinylidene difluoride (PVDF) membranes (BioRad), and analyzed with antibodies to check IP quality. Data from Proteomic Core Lab were analyzed with Scaffold software (Scaffold_5.1.2).

### Tumor models

TC1 cells, derived from mouse lung epithelial cells that were immortalized with HPV-16 E6 and E7 and transformed with the c-Ha-ras oncogene (43), were provided by Yvonne Paterson (University of Pennsylvania). Cells were grown in RPMI-1640, 10% FBS, 2 mM glutamine, and 5 μg/mL of penicillin and streptomycin. Each mouse was shaved on its right flank and injected s.c. with 1.2×10^6^ TC1 tumor cells. Tumor volume was determined by the following formula: (3.14 × long axis × short axis × short axis)/6. At 7 days after tumor injection, CHD4i was injected every day for 10 days at the concentration of 2.5 or 5 mg/kg. Tumor volume was measured every 3 days. Tumors, spleen, and LN samples were collected at the end of the experiments. Cytokines were evaluated after 4 hours induction with PMA/ionomycin. The murine hepatoma cell line H22 was purchased from Procell Life Science & Technology (Wuhan, China) and cultured as described for TC1 cells. An orthotopic tumor model was established by subcapsular injection of H22 cells (1×10^6^ in 30 μL Matrigel Basement Membrane Matrix, BD Biosciences, Franklin Lakes, NJ) into the paramedian area of the lower surface of the left liver lobe (44), with pressure using sterile swabs to prevent hemorrhage. Mice were treated daily from day 7 post-inoculation as described for the TC1 model and alpha-fetoprotein (AFP) levels were determined using mouse AFP DuoSet ELISA (#DY5369, R&D Systems, MN, USA) according to instructions of the manufacturer.

### Cardiac transplantation

Heterotopic cardiac allografts were performed using BALB/c donors and WT or Rag1^−/−^ recipients (C57BL/6 background), as described (9). In adoptive transfer studies of Treg-dependent allograft survival, after their sorting YFP^+^ Treg cells, YFP^+^ Tregs (0.5 × 10^6^) from WT or CHD4^−/−^ mice and Teff cells (1 × 10^6^) from WT mice were injected i.v. into Rag1^−/−^ mice bearing BALB/c cardiac allografts.

### Quantitative histone post-translational modification (PTM)

Quantitative histone post-translational modification (PTM) data were obtained from acid-extracted histones subjected to chemical derivatization, tryptic digestion, and targeted mass spectrometry. PTMs were quantified using Skyline as the percentage of each modified peptide relative to the total signal for the corresponding residue, enabling assessment of modification state redistribution. For each PTM, fold change was calculated as the ratio of mean relative abundance between experimental and control conditions (sgCHD4 vs sgDummy for genetic inhibition; CH41 vs DMSO for chemical inhibition). Technical replicates were averaged prior to fold-change calculation. Variability was represented by propagated standard deviation of the fold change, calculated using standard error propagation for ratios:

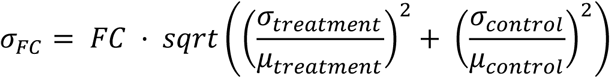

Fold changes and associated variability are visualized as dot plots with horizontal error bars, with a reference line at fold change = 1 indicating no difference relative to control. Because PTMs are expressed as relative abundance within each residue, fold changes reflect redistribution among modification states rather than absolute changes in histone abundance.

### Statistics

Data were analyzed using GraphPad Prism 10.0 and data are presented as mean ± SD unless specified otherwise. Measurements between 2 groups were done with a 2-tailed Student’s t test. Comparison of multiple samples was performed by 1-way ANOVA. Graft survival was evaluated with Kaplan-Meier followed by log-rank test. Tumor behavior after the treatment was calculated as the ratio of area under the curve (AUC) with or without the drug. P less than 0.05 was considered significant: *P < 0.05, **P < 0.01, ***P < 0.005.

### Study approval

Animal studies were approved by the Institutional Animal Care and Use Committee of the Children’s Hospital of Philadelphia (protocols IAC-22-000561 and IAC-22-001047).

## DECLARATIONS

## Acknowledgement

We thank the staff of the MS core facility at the Children’s Hospital of Philadelphia for sequencing and data analyses.

## Contributors

YX, FK and LW prepared the first draft of the manuscript. FK, MM, EdG, TA, LW undertook material preparation, data collection and analysis and, with RH and YX, carried out biochemical and animal experiments. LW carried out all other experiments. SRH and IB undertook data analysis. EN provided CHD4i and insights into CHD4 biology. WWH designed the studies, interpreted the results, and edited the final manuscript. All authors read and approved the final manuscript.

## Funding

This work was supported by an NIH grant 1R01CA253320 (to WWH), funding by the Fred and Suzanne Biesecker Pediatric Liver Center at The Children’s Hospital of Philadelphia (to WWH), and by the National Natural Science Foundation of China, Grant No. 82271810 and No. 82571041 (to YX).

## Competing interests

There are no competing interests.

## Data availability

Data are available upon reasonable request.

## Supplemental material

Additional content has been supplied by the author(s).

## Notes

**Conflict of interest**: EN and IB have filed a patent application concerning the CHD4 inhibitor used in this study. The other authors have declared that no conflict of interest exists.

### Competing Interest Statement

EN and IB have filed a patent application concerning the CHD4 inhibitor used in this study

