## Supplementary Figures & Tables for "Targeting the NuRD Component, CHD4, Impairs Foxp3+ Treg Cell Production and Function and Promotes Anti-Tumor Immunity"

[1]Institute of Hepatobiliary Diseases of Wuhan University, Transplant Center of Wuhan University, and Zhongnan Hospital of Wuhan University, Wuhan, China. [2]Department of Pathology and Laboratory Medicine, Children's Hospital of Philadelphia, Philadelphia, Pennsylvania, USA. [3]University of Udine, Udine, Italy. [4]Department of Pathology and Laboratory Medicine, University of Pennsylvania, Philadelphia, Pennsylvania, USA. [5]Cellzyme Therapeutics, Inc., San Diego, CA, USA. [6]CellarisBio, LLC, San Diego, CA, USA.

[7]These authors contributed equally: Elmar Nurmemmedov and Wayne W. Hancock.

**Conflict of interest**: EN and IB are pursuing a patent application that covers the novel CHD4 inhibitor investigated in this study. The other authors have declared that no conflict of interest exists.

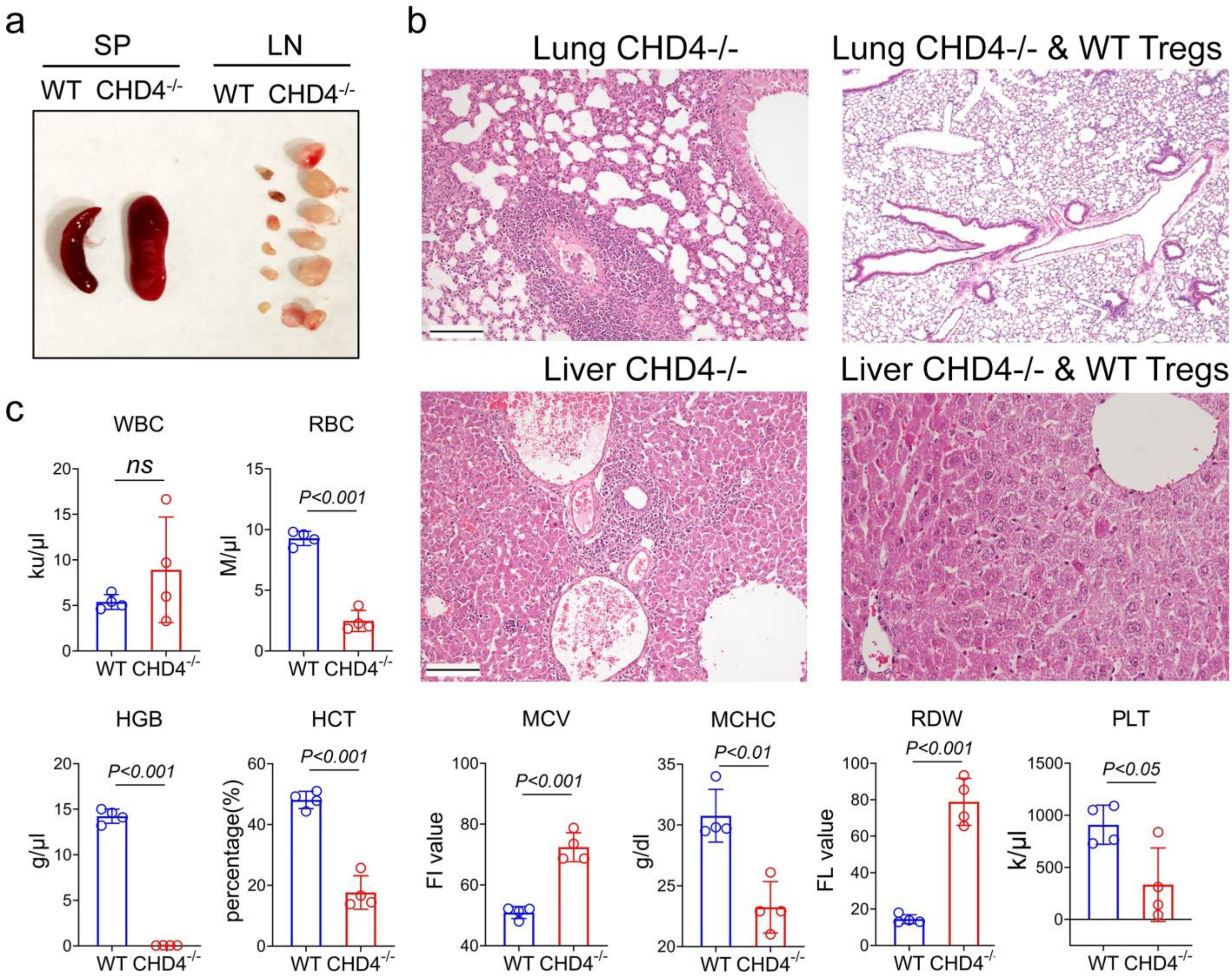


**Fig. S1. Effects of conditional deletion of *CHD4* in Foxp3$^{+}$ Tregs.** Data in panels a–c are from 20 days old mice (4 mice/group). (a) Spleen and lymph nodes from CHD4$^{-/-}$ versus WT mice. (b) Focal mononuclear infiltrates in lungs and livers of CHD4$^{-/-}$ mice at day 20 (scale bars = 100 μ) were absent in corresponding CHD4$^{-/-}$ mice rescued by i.v. injection of WT Tregs at day 3 of life. (c) Deletion of *CHD4* in Tregs was associated with decreased RBC, hematocrit (HCT), hemoglobin (HGB), mean corpuscular hemoglobin concentration (MCHC) and platelet (PLT) levels. *P < 0.05; **P < 0.01; ***P < 0.001 vs. WT control.

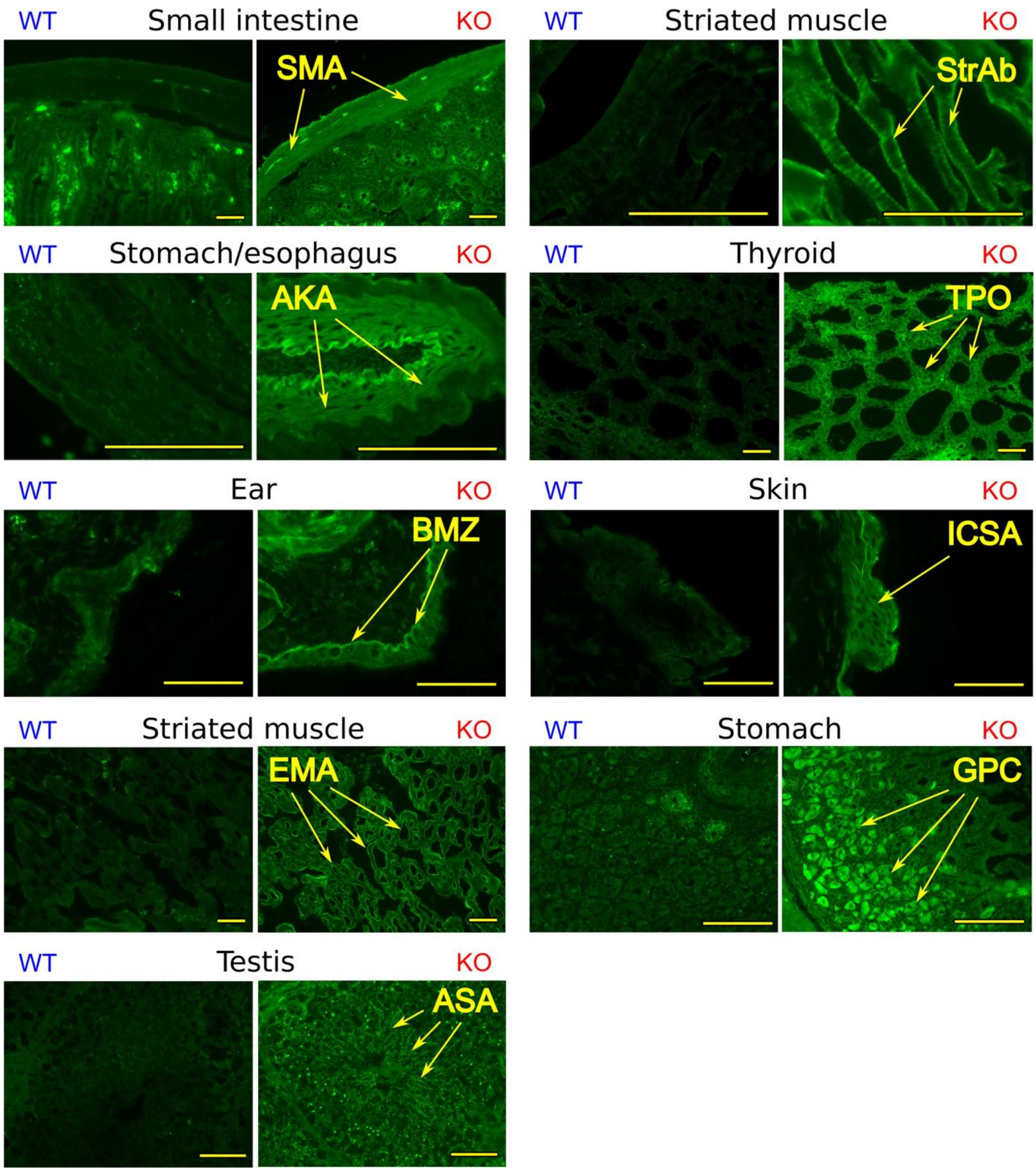


**Fig. S2. Autoantibodies detected by indirect immunofluorescence.** Sera of four CHD4$^{-/-}$ male mice 21 days old, four FOXP3-Cre control mice at the same age and two adult C57BL/6 WT mice were incubated at 1/10 dilution with cryosections of corresponding WT tissues. SMA, smooth muscle antibodies; StrAb, anti-striated muscle antibodies; AKA, anti-keratin antibodies; TPO, thyroid peroxidase antibodies; BMZ, basement membrane zone antibodies; ICSA, antibodies to intercellular cement substance; EMA, anti-endomysial antibodies; GPC, anti-gastric parietal cell antibodies; ASA, anti-sperm antibodies (reactivity to the tails of sperm cells). More data are in Supplementary Table S2.

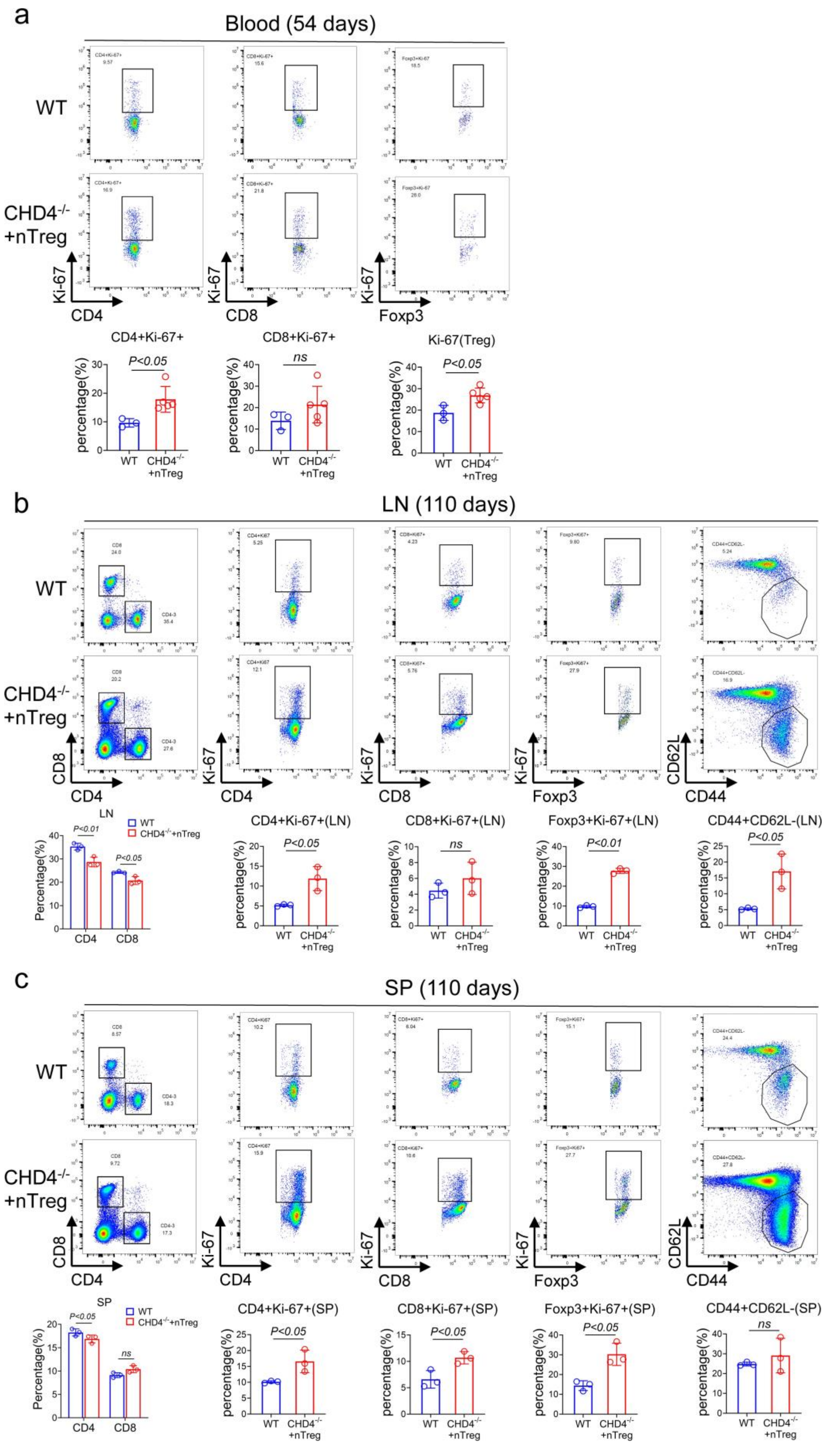


**Fig. S3. Effects of WT Treg injection on immune events in CHD4$^{-/-}$ mice.** WT Tregs ($2.5x10^6$) were injected into CHD4$^{-/-}$ mice on day 3 of life. Analysis of (a) % of Ki67 in CD8+ cells, CD4+ cells and Treg cells on day 54 after injection of WT Tregs. Analysis of the % of Ki67 in CD8+ cells, CD4+ cells, Treg cells, and CD4+CD44+CD62L$^{lo}$ cells in (b) LN and (c) spleens at 110 days after injection of WT Tregs. Data are shown as mean ± SD, 5 samples/group. Student's t-test for unpaired data; *p<0.05, **p<0.01 or ns (not significant) vs. control.

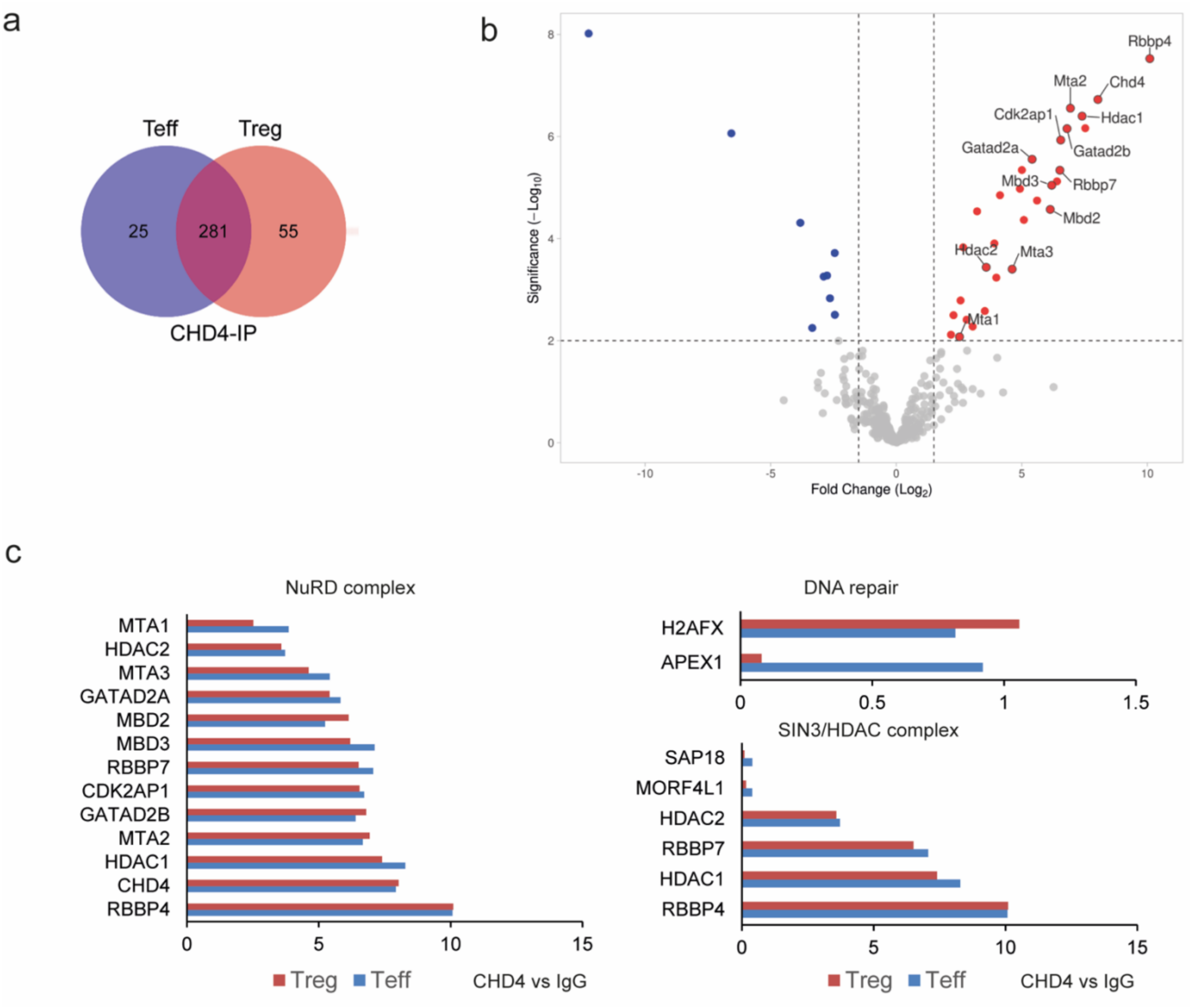


**Fig. S4. CHD4 forms highly conserved protein complexes in Treg and conventional CD4⁺ T cells. (a)** Venn diagram of CHD4-interacting proteins identified by CHD4 immunoprecipitation followed by mass spectrometry in Treg and Teff cells. **(b)**Differential enrichment analysis of CHD4-associated proteins in Treg versus Teff cells. Core NuRD components are highlighted. **(c)** Normalized spectral counts of representative CHD4 interactors belonging to the NuRD, SIN3/HDAC and DNA repair complexes, demonstrating a largely conserved CHD4 interactome between Treg and Teff cells. Values are shown as CHD4 versus IgG enrichment after normalization for total spectra.

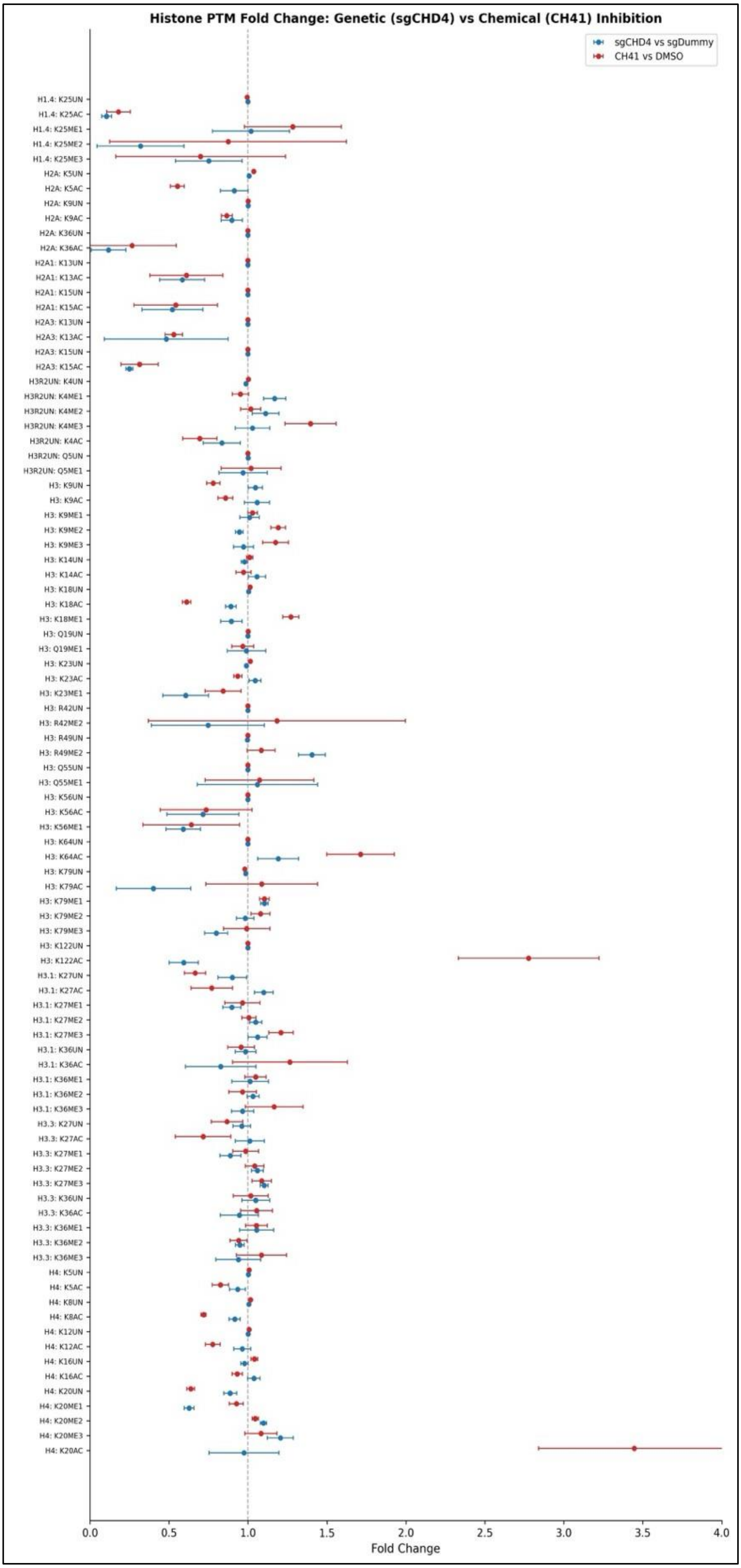


**Fig. S5. Comparative histone PTM redistribution following genetic vs. chemical inhibition of CHD4 in human NSCLC cells.** Fold changes in histone PTM abundance are shown for genetic CHD4 depletion (sgCHD4 vs sgDummy, blue) and chemical CHD4 inhibition (CH41 vs DMSO, red). Histones were acid-extracted, chemically derivatized, digested with trypsin, and analyzed by targeted mass spectrometry, with PTMs quantified as the percentage of each modified peptide relative to the total signal for the corresponding residue. Points represent mean fold change for each PTM, and horizontal error bars indicate standard deviation derived from technical replicates. The vertical dashed line at 1 denotes no change relative to control.

**UN** – Unmodified

**ME1/2/3** - Mono/di/tri methylation

**AC** – Acetylation

**UB** – Ubiquitination

**K27M** - Mutation of lysine to methionine

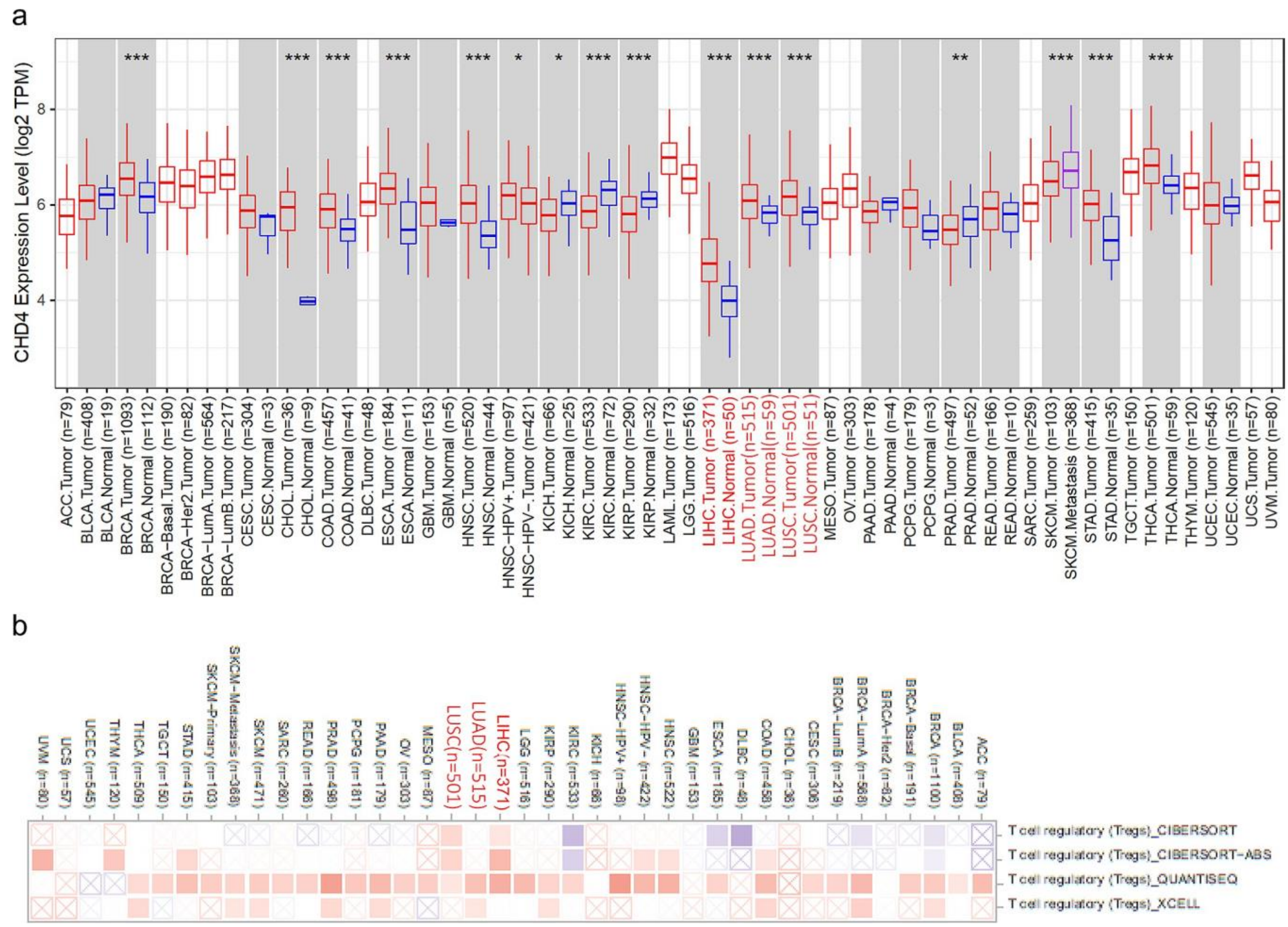


**Fig. S6. CHD4 is essential for Treg biology in NSCLS.** (a) Differential expression of CHD4 in tumors vs. adjacent normal tissues (all TCGA tumors). (b) Correlation between CHD4 expression and Treg infiltration in cancers.

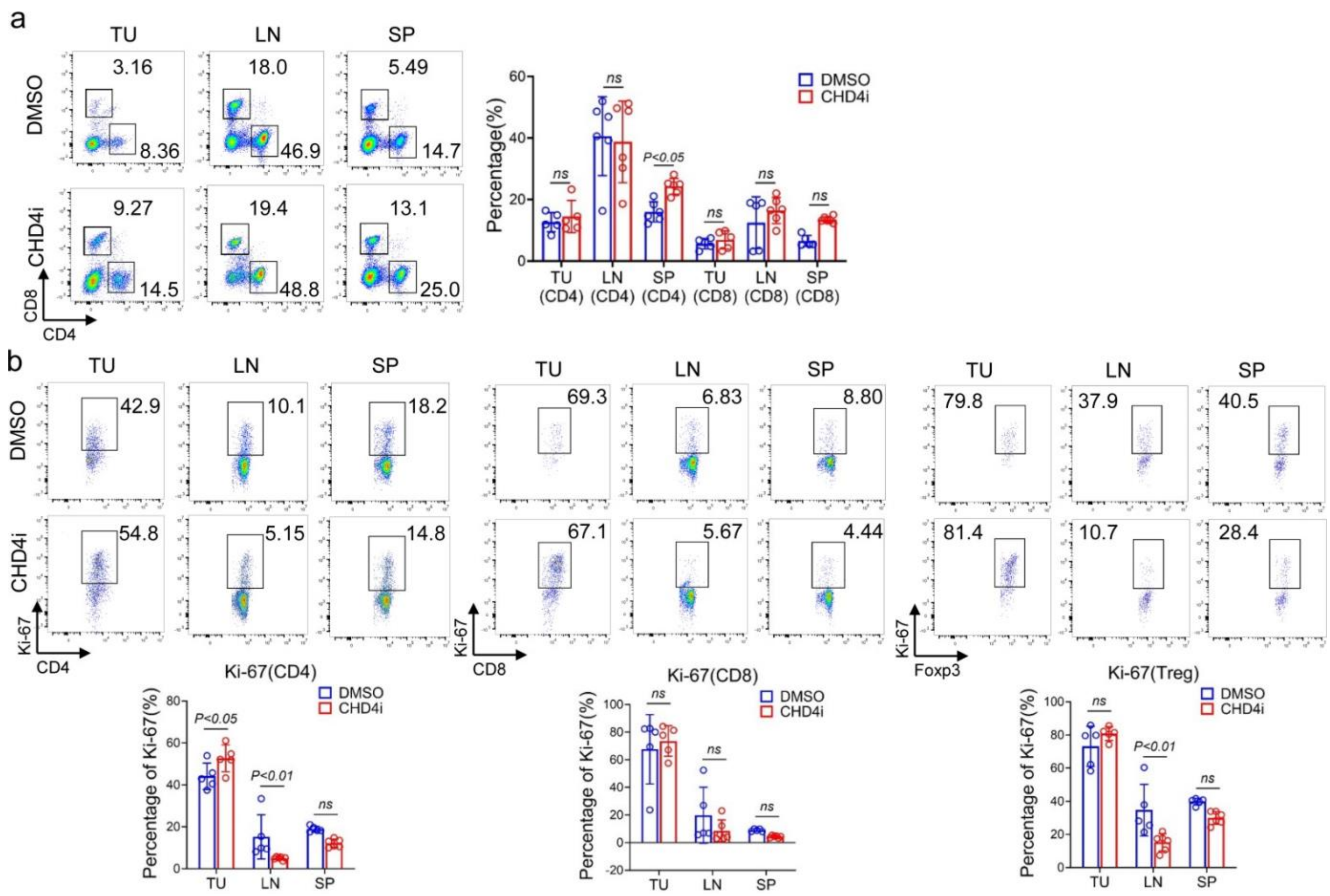


**Fig. S7. CHD4i enhances anti-tumor immunity (HCC model).** Analysis of the percentages of (a) CD4+, CD8+, CD4+Ki67+, CD8+Ki67+, and Foxp3+Ki67+ cells in lymph nodes, spleens and tumors from CHD4i and DMSO-treated mice. Data are shown as mean ± SD, 8-10 samples/group. Student's t-test for unpaired data; *p<0.05, **p<0.01 or ns (not significant)

**Supplementary Table 1**

Histopathology of i) male mice at 15 days of age with conditional deletion of *CHD4* in Tregs and ii) CHD4-/- male mice treated with normal Tregs (nTregs) and followed to >110 days of age[1]

| Tissue | Histologic findings (CHD4 KO) | Histologic findings (CHD4 KO + nTregs) |
|---|---|---|
| Lung | Widespread MNC infiltrates, especially adjacent to bronchovascular bundles. | Minor residual infiltrates primarily centered on large bronchovascular bundles |
| Liver | Periportal MNC infiltrates and piecemeal hepatocyte necrosis. | Liver lobe structures are intact, and there is no lymphocyte infiltration in the portal area, central vein, hepatocytes or liver sinusoids. |
| Spleen | Splenic parenchyma is normal with well-organized peri-arteriolar lymphoid sheaths and lymphoid follicles. Scattered foci of extramedullary hematopoiesis are identified within the intact red pulp. | Normal histology |
| Lymph nodes | Nodes display normal cellularity but there is effacement of nodal architecture, with loss of distinct follicles (inconspicuous germinal center formation). | Normal histology |
| Thymus | Overall thymic architecture is intact with preservation of distinct corticoid-medullary junctions. No apparent decrease in the volume of cortical thymocytes was seen. | Normal histology |
| Skin | A diffuse, mild MNC infiltrate predominantly present within the superficial dermis, with focal exocytosis. | Overall skin architecture is intact and there is no MNC infiltration of hair follicles and small blood vessels. |
| Pancreas | Pancreata are normally developed with well-formed islets of Langerhans. Rarely, perivascular inflammation was noted in peri-pancreatic adipose tissue. | Normal histology |
| Heart | Unremarkable cardiac myocytes; rare foci of extramedullary hematopoiesis. | Normal histology |
| Kidney | Renal parenchyma, including glomeruli and tubules, was normally developed. No significant interstitial or perivascular inflammation. | Normal histology |
| Gastro -intestinal tract | Small bowel contains scattered collections of MNC in the lamina propria without infiltration into surface epithelium. Villous architecture is intact. | Small bowel has scattered collections of MNC in the lamina propria without infiltration into surface epithelium. Villous architecture is intact. |

[1]Tissues from 4 mice/group were fixed in 10% neutral buffered formalin, routinely processed, embedded in paraffin and stained with hematoxylin and eosin.

**Supplementary Table 2**

Autoantibodies in mice with conditional CHD4 deletion in Foxp3+ Treg cells
and effects of early Treg adoptive cell transfer

| N | Type of autoantibodies | Substrates | Present at 21st day, in untreated mice | Treg-treated mice | | |
|---|---|---|---|---|---|---|
| | | | | Day 42 | Day 54* | Day 110 |
| 1 | Anti-nuclear (ANA) | Liver, kidney and all other tissues | No | No | No | No** |
| 2 | Anti-mitochondrial (AMA) | Liver, kidney, stomach | No | No | No | No |
| 3 | Anti-smooth muscle (SMA) | Colon, small intestine, stomach | Yes, 3 of 4 | No | No | Yes, 3 of 5 |
| 4 | Anti-striated muscle (StrAb) | Striated muscle | Yes, 4 of 4 | No | No | No |
| 5 | Anti-keratin (AKA) | Skin, ear, esophagus | Yes, 4 of 4 | Yes, low, 3 of 5 | No | Yes, 3 of 5 |
| 6 | Anti-islet (ICA) | Pancreas | No | No | No | No |
| 7 | Anti-steroid-producing cells | Adrenal gland, testis | No | No | No | Yes, 1 of 5 |
| 8 | Anti-thyroperoxidase (TPO) | Thyroid gland | Yes, low, 2 of 4 | No | Yes, low, 2 out of 5 | Yes, low, 2 of 5 |
| 9 | To epithelial basement membrane zone (BMZ) | Skin, ear | Yes, 4 of 4 | Yes, low, 2 of 5 | No | No |
| 10 | To intercellular cement substance (ICSA) | Skin, ear | Yes, 3 of 4 | No | Yes, low, 5 out of 5 | Yes, 3 of 5 |
| 11 | Anti-endomysium (EMA) | Striated muscle | Yes, 2 of 4 | No | No | Yes, 5 of 5 |
| 12 | Anti-gastric parietal cell (GPC) | Stomach | Yes, 3 of 4 | Yes, 1 of 5, very high | Yes, 1 out of 5, high | Yes, 1 of 5, very high |
| 13 | Anti-sperm | Testis | Yes, 3 of 4 | Yes, low, 1 of 5 | No | No** |

* Due to small volumes, sera samples were combined as mouse 1+2 and mouse 3+4+5, therefore results are presented for the group of mice.

** At day 110 post-Treg transfer, some unusual and not previously described autoreactivities patterns were found, such as anti-nuclear staining exclusively in striated muscle cells, reactivity to sperm acrosomal antigen, reactivity to spermatogonia cells, tight junction reactivity in liver and pancreas, and others.
